# The stability of fatty acid composition in sunflower oil is dependent on environment and affected by structural variation

**DOI:** 10.64898/2026.05.04.722759

**Authors:** Markus Ingold, Qingming Gao, Jennifer R. Mandel, James P. McNellie, Kyle G. Keepers, Jessica G. Barb, John M. Burke, Loren H. Rieseberg, Brent S. Hulke

## Abstract

In sunflower (*Helianthus annuus* L.), the composition of fatty acids in the seeds, primarily oleic, linoleic, stearic and palmitic acid, is of utmost importance for oil quality. Despite this, the genetic basis of this trait and its interaction with the environment is poorly understood. Understanding this interaction is critical to improvement of sunflower within the context of climate change. In this work, we incorporated fatty acid composition measurements from the sunflower SAM population and eight environments across an extensive geographic cline into GWAS. The SAM panel consists of 287 varieties representing approximately 90% of sunflower diversity, for which 2.2 million high-quality SNPs with a MAF > 5% are available. For increased power, multivariate GWAS was performed with four different inputs: (i) mean fatty acid composition within each environment, (ii) mean fatty acid composition within each environment omitting high oleic varieties, (iii) trait stability within environments quantified by standard errors among replicate samples (α stability) and (iv) Eberhart and Russell’s β which quantifies trait stabilities across environments (β stability). All four analyses yielded highly significantly associated SNPs. We found that high oleic varieties exhibited high β trait stability, resulting in substantial overlap in markers between analyses (i) and (iv), with signals being fairly consistent between environments in analysis (i). For analyses (ii) and (iii), significant markers tended to vary between trials. For significant SNPs across all analyses, 147 candidate genes were identified, including promising candidates such as 15 fatty acid metabolism genes, 6 heat shock proteins and 22 transcription factors. Lastly, a large introgression consisting of two flanking inverted sequences on Chromosome 5 was found to coincide with α stability in the Georgia trial, suggesting a role in FA composition stability under high heat conditions.

## INTRODUCTION

Sunflower (*Helianthus annuus* L.) was originally domesticated from wild progenitors in North America, ca. 4000 years ago (Crites, 1993). After being introduced to Europe in the 1500s, domestication continued, and varieties were bred for seed oil content, increasing from 33% in 1913 to 51% in 1958 during selective breeding in the former Soviet Union (Enrique et al., 2015). The high oleic (HO) genotype was obtained in the same region by inducing mutations using dimethylsulphonate on seeds, resulting in the Pervenets variety (Soldatov, 1976). While individual seeds of this variety were reported to vary between 190 and 940 g/kg of oleic acid across environments, the HO composition of its offspring was exceedingly stable (Urie, 1985). The source of this HO phenotype is thought to be in large part caused by a partial duplication of the seed-specific *fatty acid desaturase 2-1* (*FAD2-1*) gene (Schuppert et al., 2006), silencing its expression and thereby inhibiting its function of catalyzing a reaction converting oleic acid to linoleic acid (Dar et al., 2017). Presently, sunflower is an important oilseed crop, grown globally for its high-quality oil (Škorić, 1992), its capability to reliably produce seed with high compositions of oleic acid (up to 930 g/kg oleic acid; Hulke and Kleingartner, 2014) and its relatively high resistance to heat and drought (Tahir et al., 2002).

The fatty acid (FA) composition of sunflower oil has a significant impact on its industrial and nutritional value, and this complex trait is influenced by both environmental and genetic factors and their interactions with each other (Rauf et al., 2017). However, few genome-wide association studies (GWAS) or quantitative trait locus (QTL) analyses have been performed to elucidate genetic factors influencing fatty acid composition in the seed (Chernova et al., 2021; Ebrahimi et al., 2008; Pérez-Vich et al., 2002; Pérez-Vich et al., 2016), and to the authors’ knowledge, no multivariate GWAS involving multiple fatty acids simultaneously has been performed in sunflower. Further, no study has investigated the genetics of FA phenotype stability over a gradient of climates, in order to develop breeding goals in preparation for climate change or for new environments, more generally. One additional issue is the instability across generations of the HO phenotype in certain varieties, the origin of which is unknown but is a recurring issue for breeders. The present study aims to fill these knowledge gaps by performing GWAS, genotype-by-environment, and candidate gene analyses on the sunflower association mapping (SAM) population, 287 varieties that capture 90% of sunflower diversity (Mandel et al., 2011), grown in 8 different location-time combinations.

Many measures of phenotypic stability have been proposed over time, two of which are potentially useful and easily applied to study instability of the phenotype and genotype-by-environment effects on the fatty acid phenotypes. Within-environment stability of a variety can be simply quantified as the phenotypic standard error of replicates; however, among-environment stability is often conveyed by the regression slope of the rankwise-sorted grand means of phenotype values in each environment (β statistic as described by Eberhart and Russell, 1966). Fatty acid composition stability within and between environments is of economic interest, as fluctuations in oil quality for the same plant variety across environments lead to inconsistent quality of the crop, potentially reducing its value. It has been shown that the fatty acid composition in sunflower becomes increasingly unstable with rising heat and drought stress, with a tendency to decrease levels of polyunsaturated FA (linoleic acid; Schulte et al., 2013; Rondanini et al., 2003). Previously, GWAS was used to discover loci associated with abiotic stress resistance in sunflower, identifying QTLs significantly associated with the plasticity of total oil yield under water-, nitrogen-, salt-, heat- and cold-stress (Mangin et al., 2017; McNellie et al., 2024). In contrast, the present study focuses on the genetics of FA variability among varieties and FA composition stability within and among environments. To understand the genetics and facilitate breeding of varieties that retain desirable FA compositions in different environments, what we call α stability (standard errors of fatty acid composition within an environment) and β stability among the 8 trials were investigated in addition to mean FA composition using multivariate GWAS.

## METHODS

### 2.1 GENOTYPE AND PHENOTYPE DEVELOPMENT

The SAM population was grown in eight field trials in North America: Vancouver, British Columbia (2010); Moorhead, Minnesota (2015, early and late plantings 2016); Ames, Iowa (2010, 2013 and 2014); and Athens, Georgia (2010). The experimental design in each environment was a randomized, complete block design with 2 to 4 plants sampled within each 6.1 m-long plot. Plants were covered with Delnet head bags prior to bloom and harvested individually. After threshing, 20 seeds were collected from each head and individually subjected to gas chromatography using either a HP 5890 or an Agilent 8890 gas chromatograph, and a fatty acid methyl ester protocol (Hulke et al., 2010). Means and standard errors were calculated for palmitic, stearic, oleic, and linoleic acid in each sunflower line in each environment. Eberhart and Russell’s β stability was calculated from means across environments (Eberhart and Russell, 1966), with genotypes absent in more than three trials or missing in the Georgia trial (i.e., a humid, subtropical outlier environment) omitted to avoid skewing.

SNP markers were retrieved from http://www.helianthome.org/download/#genotype (Bercovitch et al., 2022). These SNP markers were called from the HA 412HOv2.0 reference genome (Huang et al., 2023) using GATK best practices (Auwera et al., 2013; Auwera and O’Connor, 2020). The resulting VCF was filtered for single copy sites based on depth. Additionally, filtering was done to select sites with a minimum quality score of 100 (minQ=100), minor allele frequency of 0.05 or greater (MAF≥0.05) and max missingness value of 0.9 (max-missing=0.9). Missing data were imputed using BEAGLE version 5.3 with default settings retained (Browning et al., 2018). This filtering resulted in a total of 2.2 million high-quality SNPs. The VCF was converted to PLINK format using PLINK beta (v1.90b6.27; Chang et al., 2015).

### 2.2 CLIMATE DATA ASSOCIATION

For each growing environment, the climate moisture index, near-surface relative humidity, potential evapotranspiration, precipitation amount, surface downwelling shortwave flux in air, near-surface wind speed, mean daily air temperature, mean daily maximum air temperature, mean daily minimum air temperature and vapor pressure deficit were extracted from the CHELSA dataset (accessed 20 April 2026; Karger et al., 2017a; Karger et al., 2017b) for the month of planting and the two subsequent months using the R library raster3 (v3.6-14; Hijmans, 2023). Iowa 2010 was omitted due to a larger number of missing samples. The Pearson correlations between the climate variables, in addition to latitude and longitude, and mean fatty acid compositions and standard errors (α stability) were then calculated and visualized. GGE biplots were created for both FA compositions and α stability using the R package gge (v1.7; Wright and Laffont, 2021).

### 2.3 UNIVARIATE AND MULTIVARIATE GWAS

ADMIXTURE (v1.3.0; Alexander et al., 2009) was used on the filtered VCF. As no evidence for population structure was found, the kinship matrix created using GEMMA (v0.98.5; Zhou and Stephens, 2012) was deemed sufficient. Univariate and multivariate GWAS was performed for the FA compositions, FA standard errors of each environment independently (termed within-environment α stability in this work), and FA β stability. For the multivariate FA compositions and β stability, GWAS was additionally performed under omission of high oleic varieties, determined by the presence of the *FAD2-1* mutation found in Pervenets (Table S1). A univariate analysis was also performed using the *FAD2-1* mutation as a covariate for each fatty acid. Univariate GWAS were performed using vcf2gwas (v0.8.7; Vogt et al., 2022), and 4 fatty acid multivariate GWAS were performed using GEMMA (v0.98.5; Zhou and Stevens, 2012).

### 2.4 POST-GWAS ANALYSIS

GWAS results were visualized using the R packages qqman (v0.1.8; Turner, 2018) and CMplot (v4.2.0; LiLin, 2022) for Manhattan and qq plots. Bonferroni correction on the number of SNPs and the number of LD blocks were used to calculate the QTL significance thresholds. The number of LD blocks was computed using LDBlockShow (v1.40; Dong et al., 2021) on a VCF pruned for an LD distance of 10,000 base-pairs.

The four linoleic acid LD blocks and four oleic acid LD blocks from Chernova et al. 2021 Table 1 were subset as fastas from the HanXRQr2.0-SUNRISE-2.1 genome (Badouin et al., 2017) using samtools faidx (Li et al. 2009). We note that two typos in the table were corrected. The linoleic-associated block on chromosome 11 from 5,004,818 to 50,619,247 was corrected to occur from 50,204,828 to 50,619,247. Additionally, the linoleic-associated LD block on chromosome 11 from 95,051,157 to 92,468,132 was assumed to occur from either 92,051,157 to 92,468,132 or 95,051,157 to 95,468,132. Both of these latter regions were included in the homology search against HA 412HOv2.0. The regions were queried in a blast search against the HA 412HOv2.0 genomic assembly using blastn (Camacho et al. 2009). The resulting tables were sorted to only include the hits with the top 20 largest bitscores, and homologous regions in HA 412HOv2.0 were identified on the basis of (1) belonging to the same ordinal chromosome, (2) if split among multiple blast hits, occurring over a sequential region that corresponded to a highly similar region of the HanXRQr2.0 chromosome, and (3) exhibiting synteny across the entirety of the LD block and the HA 412HOv2.0 assembly.

A list of genes expressed in developing sunflower seeds was assembled to only include genes with a log2 fold-change > 2 between developing seed and 10 other tissues (Badouin et al., 2017). This gene list was further reduced to only include entries with a protein function predicted by MapMan (X4.2_helianthus_annuus.txt from mapman.gabipd.org; Thimm et al., 2004) and those around significant SNPs that appeared in at least 4 GWAS results, with a maximum distance of 1 Mb from each locus peak. The GWAS results that were compared included multivariate mean FAs with and without *FAD2-1* mutants in each environment, multivariate α stability in each environment, and multivariate β stability. A Venn Diagram comparing these results was created using the R package VennDiagram (v1.6.20; Chen, 2018).

## RESULTS

### 3.1 PHENOTYPE ANALYSIS

The SAM population was grown in 8 environments and the seed oil composition of the four major FAs was measured. In Figure S1, the relationships between these FAs are displayed, showing a major negative correlation between oleic and linoleic acid compositions. This is expected because of the known tradeoff between oleic and linoleic acid caused by different alleles at the *FAD2-1* gene. While the correlations between the rest of the FAs were smaller, all correlations except between palmitic and stearic acid were statistically significant, highlighting their physiological relationships.

Eighty-one of the varieties were grown in the seven environments for which weather data was available (Iowa 2010 is excluded due to missing samples). Only one of these varieties was of the HO phenotype. The FA compositions in different environments were correlated to several available climate variables in the month of planting and the two subsequent months (Figure 1). Oleic and linoleic acid compositions show significant correlations with latitude, vapor pressure deficit, mean daily minimum (night temperature) and maximum temperature, and mean daily temperature during the third month after planting, which coincides with the time of seed development. Additionally, oleic acid shows a significant correlation with longitude. Stearic acid shows the largest correlations with climate variables during the first month after planting, with mean temperature and cloud cover reaching the significance threshold.

**Figure 1:**
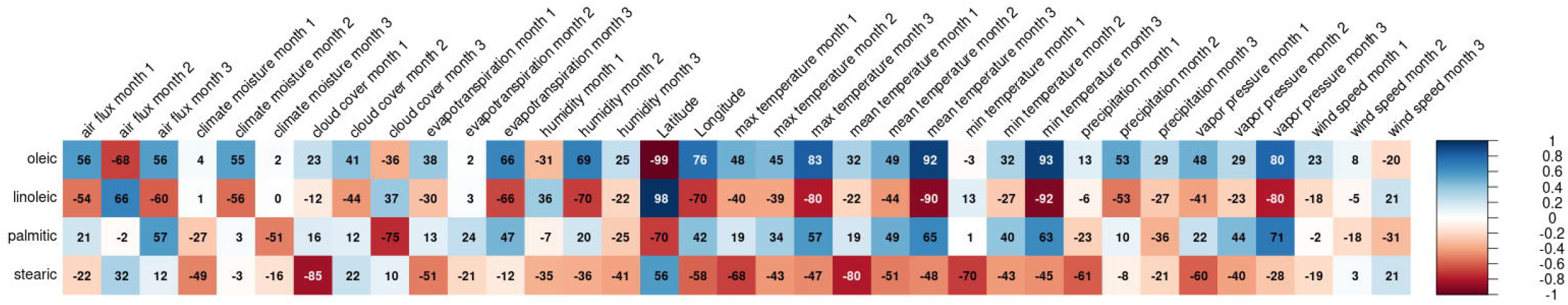
Correlations between the mean FA compositions of 81 sunflower lines present in 7 environments and various climate variables. Monthly climate variables were extracted from the CHELSA dataset, with month 1 signifying the month of planting. Positive correlations between FA compositions and climate variables are marked in blue, negative ones in red, with color intensity denoting correlation strength. Significant correlations (p < 0.05, without p-value correction) are marked by white text.

Additionally, a genotype plus genotype-vs-environment interaction (GGE) analysis was conducted and visualized as a biplot in Figure 2 (see also Tables S2 and S3). In it, the tradeoff relationship between oleic and linoleic acid is consistently observed, similar to Figure S1, but the tradeoff between oleic and palmitic acid is more pronounced when all varieties are included. This latter tradeoff largely disappears when high oleic (*FAD2-1* mutant) lines are removed, instead showing a direct tradeoff between stearic and palmitic (the two major saturated fatty acids) and oleic and linoleic (the two major unsaturated fatty acids). Furthermore, environmental interactions of stearic acid seem to be considerably different from the other three FAs, as was observed in Figure 1. High oleic and relatively high stearic varieties are pronounced in their separation from the majority of samples in this plot.

**Figure 2:**
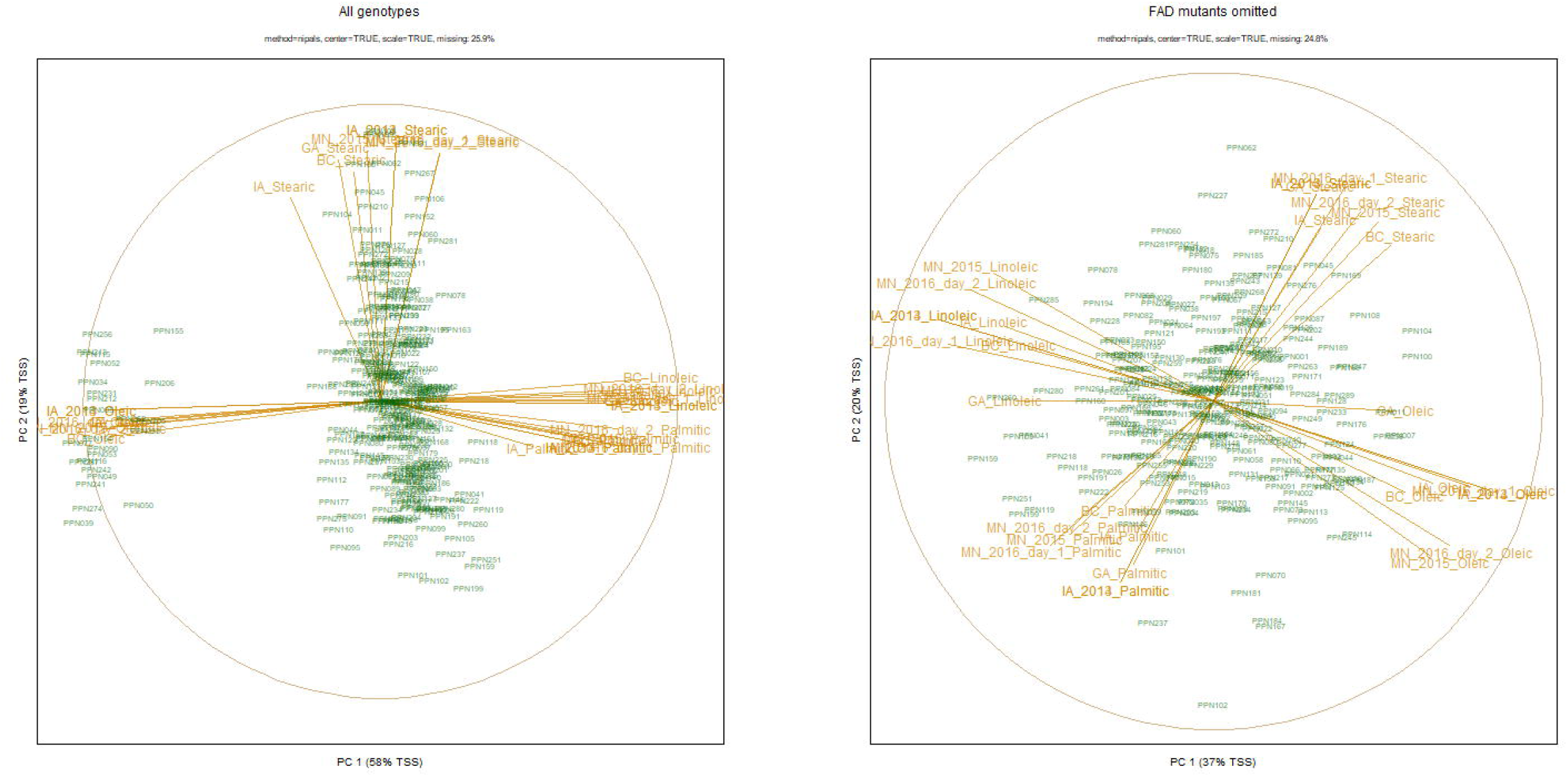
Genotype plus genotype-vs-environment interaction biplots. The left biplot includes all varieties, the right biplot excludes variants with the *FAD2-1* mutation. In the left biplot, the cluster on the left side contains all of the high oleic sunflower lines, with the center cluster showing a continuum from top to bottom of the highest stearic acid content to the lowest stearic content. In the right biplot, variation exists for all four major fatty acids, with strong tradeoffs between the two major saturated fatty acids (palmitic and stearic) and the two major unsaturated fatty acids (oleic and linoleic). Gold colored vectors are fatty acid-environment combinations and green points are sunflower lines. Exact vector loadings can be obtained from Supplemental Tables S2 and S3.

The β stability of component fatty acids in varieties varies considerably, from highly stable (index near or below zero) to low stability (index greater than four) in the case of palmitic acid (Figure 3). When FAD mutants and the WT are compared, the HO varieties had significantly higher β stability for oleic and linoleic acid composition, with no significant difference for stearic and palmitic acid.

**Figure 3:**
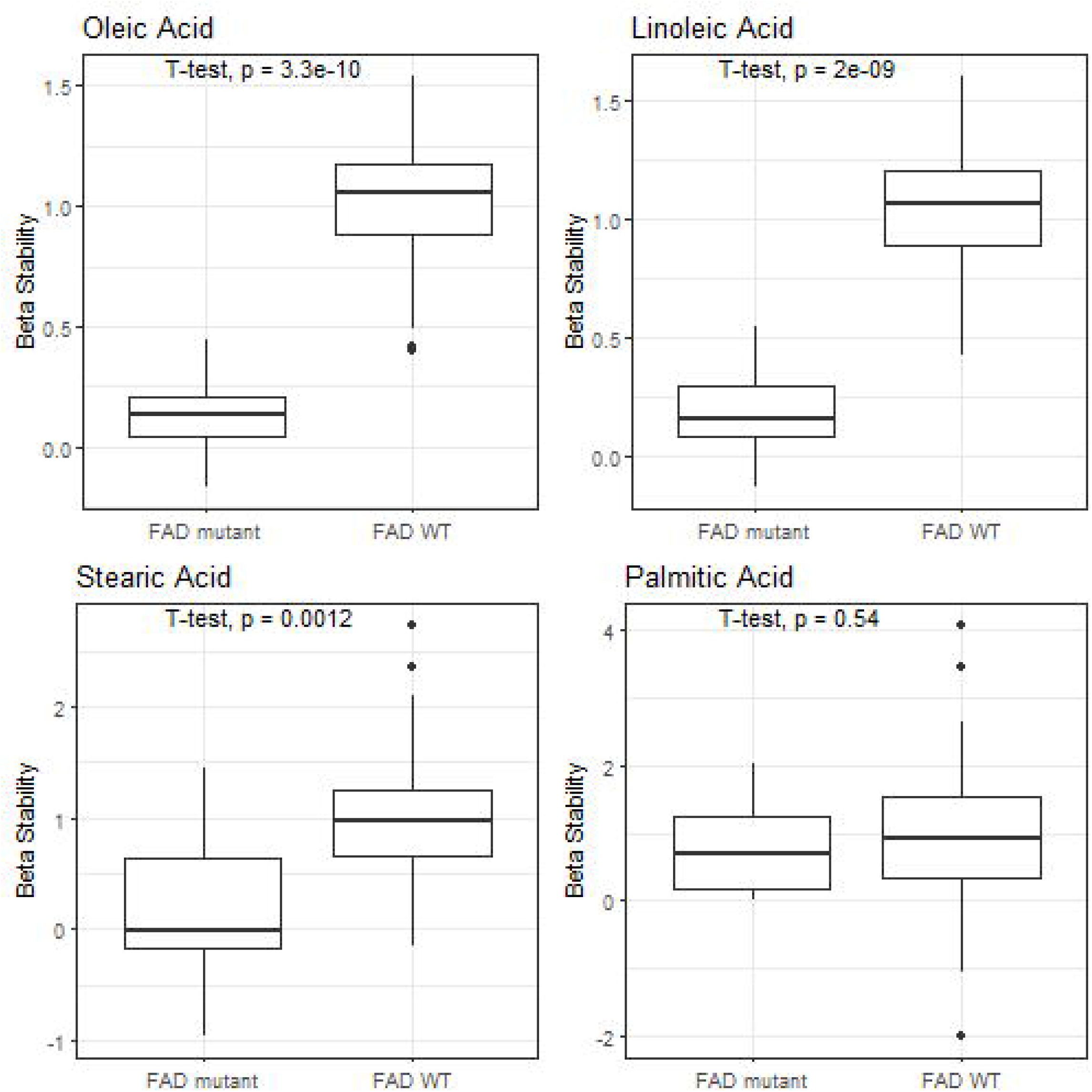
Boxplots of β stability of the four major fatty acids, grouped by the presence of the *FAD2-1 mutation*. Lower values signify higher trait stability.

### 3.2 GWAS ON MEAN FATTY ACID COMPOSITION

GWAS on the SAM population yielded highly significant associations in every environment, with −log_10_(p) reaching more than 40. While the highest peak on chromosome 14 coincides with the haploblock on which the *FAD2-1* mutation is located, multiple significant peaks on other chromosomes are present. As seen in Figure 4, peaks on chromosomes 1, 11, 13, 14 and 16 are supported by multiple environments (see also Table S4).

**Figure 4:**
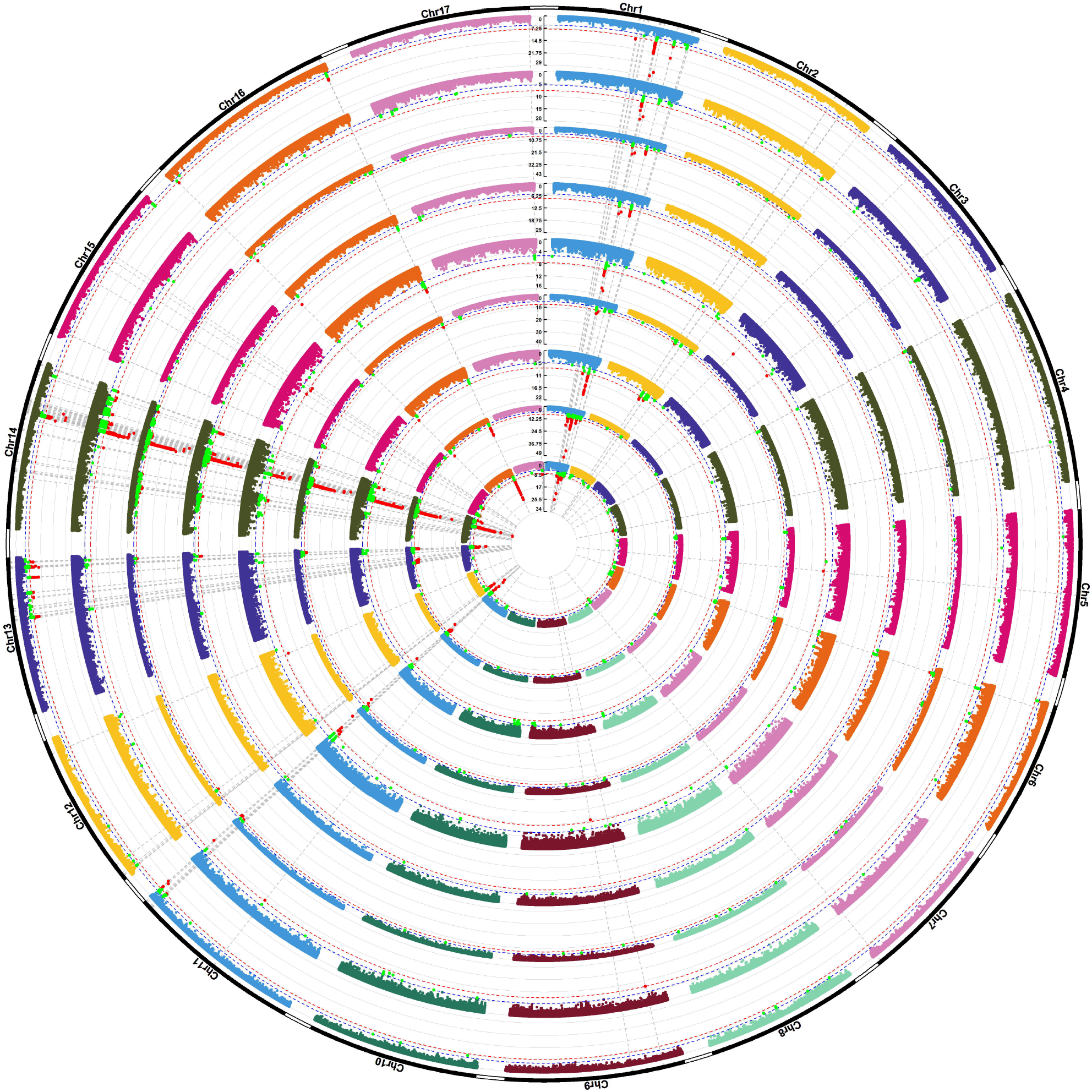
Manhattan plots of the multivariate response for mean fatty acid composition in all trials vs. the *β* stability. From inside to outside, the Manhattan plots represent the results of the trials in British Columbia 2010; Minnesota 2015 and early and late 2016; Iowa 2010, 2013 and 2014; Georgia 2010; and Eberhart and Russell’s *β* stability. SNP positions are depicted on the x-axis, −log10 of the p-value on the y-axis. The Bonferroni threshold based on the number of SNPs is depicted as a red line, the Bonferroni threshold based on the number of LD blocks as a blue line. SNPs passing the lower threshold are marked green, SNPs passing the more stringent threshold are marked in red.

### 3.3 GWAS ON TRAIT STABILITY ACROSS ENVIRONMENTS

Using Eberhart & Russell’s β, the stability of the four major FAs was quantified and used as input for multivariate GWAS. The resulting Manhattan plot in Figure S2 shows significant associations on chromosomes 1, 11, 13, 14 and 16, with the highest peak on chromosome 1. Strikingly, the QTLs for β stability coincide with QTLs for FA composition, as shown in Figure 4 (see also Table S4). Plotted alongside the mean FA composition GWAS in Figure 4, it becomes clear that the majority of peak locations coincide between said analyses and β stability. Interestingly, on chromosome 13, stability-associated SNPs are distributed along a larger, contiguous region when compared to associations with the mean multivariate fatty acid composition at each of the 8 environments.

### 3.4 OMISSION OF HIGH OLEIC VARIETIES IN GWAS

To detect smaller effects that are potentially masked by the major locus for high oleic composition, all 28 samples with the causal *FAD2-1* duplication were omitted from the following GWAS. Figure S3 depicts the Manhattan plots of these analyses (see also Table S4). Compared to Figure 4, while significant SNPs are still present, p-values are much less extreme. Further, far fewer QTLs are supported by multiple trials. Those with multi-environment support are present in at most four environments (e.g. chromosome 6). The β stability GWAS does not reach the strict Bonferroni threshold after HO varieties are omitted (not shown).

The difference in results from using all vs. omitting HO varieties becomes clear in the qq plot in Figure S4, comparing p-values between the two analyses. Omitting HO varieties leads to weaker associations in line with the less extreme remaining phenotypes; however, an abundance of very low p-values appears when HO varieties with the *FAD2-1* mutation are included. This result would traditionally flag caution in the GWAS analysis, except that a large number of SNPs are associated with the *FAD2-1* mutation due to high linkage disequilibrium in the region. The high linkage disequilibrium is due to the rapid conversion of genetic materials to the HO phenotype in recent years. To show that the p-values are not an artifact of the multivariate approach, qq plots for univariate GWAS on oleic acid were additionally visualized, with results similar between the univariate and multivariate approaches. Univariate analyses for each FA and environment for both the full dataset and the dataset omitting HO varieties are shown in Figures S6 to S13.

### 3.5 GWAS ON TRAIT α STABILITY WITHIN ENVIRONMENTS

For the analysis of α stability, Iowa 2010 and Minnesota 2015 were omitted due to missing replicates. For the remaining trials, significant associations exist, as visualized in Figure 5 (see also Table S4). Even though highly significant peaks are visible on all chromosomes except 4, few of these QTLs are supported by multiple environments in the multivariate analysis. Overlaps of significant SNPs between environments are visible on Chromosomes 5, 7 and 13. Associations on these chromosomes are also present in univariate analyses, particularly for oleic and linoleic (Figure S14 to S17).

**Figure 5:**
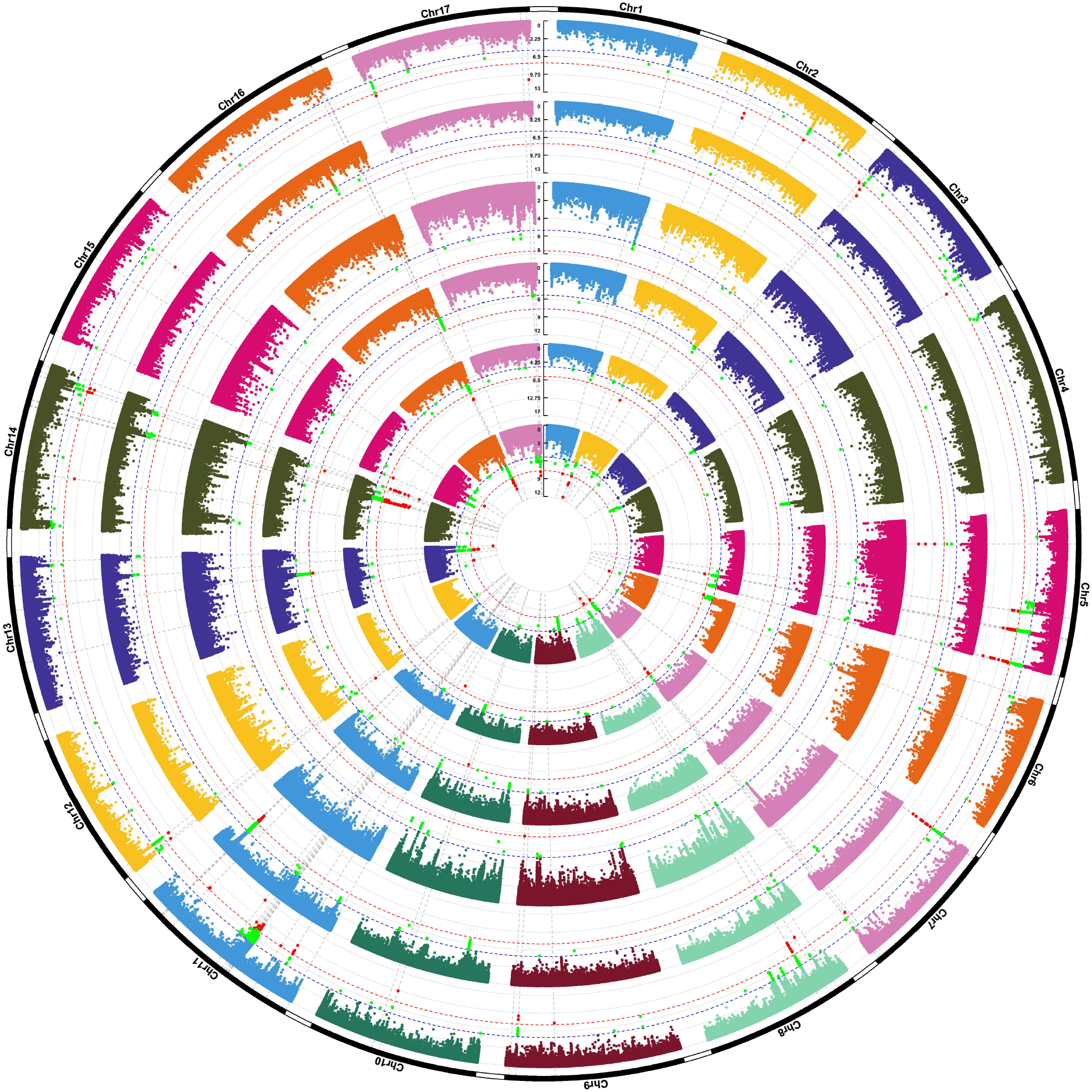
Manhattan plots of multivariate fatty acid composition standard errors (α stability) in all six trials with available data. From inside to outside, the Manhattan plots represent the results of the trials in British Columbia 2010, Minnesota early and late 2016, Iowa 2013 and 2014 and Georgia 2010. SNP positions are depicted on the x-axis, −log10 of the p-value on the y-axis. The Bonferroni threshold based on the number of SNPs is depicted as a red line, the Bonferroni threshold based on the number of LD blocks as a blue line. SNPs passing the lower threshold are marked green, SNPs passing the more stringent threshold are marked in red.

### 3.6 CANDIDATE GENE ANALYSIS

For the candidate gene analysis, all expressed genes were first subset by high expression in the developing seed compared to other tissues. Genes of this subset flanking significant SNPs within 1 million basepairs were then identified as candidate genes. A total of 64 candidates were identified for the mean FA analysis without FAD omission (Figure 4, Table S5), 68 for the mean FA analysis with FAD omission (Figure S3, Table S5), 22 for the β stability analysis (Figure S2, Table S5) and 58 for the α stability analysis (Figure 5, Table S5). As shown in Figure S5, several genes were found in multiple analyses, with substantial overlap between mean FA with and without FAD mutants as well as β stability. Only relatively few genes identified for standard errors overlap with the other analyses, with a notable lack of genes shared with the β stability.

## DISCUSSION

### 4.1 PHENOTYPE ANALYSIS

The field trials used in this work span from a subtropical climate in Georgia to moderate oceanic temperate rainforest in British Columbia. Climate variables at the same locations in Iowa and Minnesota vary by year, and in addition, early and late planting in Minnesota in 2016 capture hotter and cooler growth conditions, resulting in 8 environments with a range of various stresses. The trend observed by Schulte et al. (2013), in which heat stress reduced the linoleic acid to oleic acid ratio in sunflower, was also observed in the present study (Figure 1). The mean daily minimum temperature, or night temperature, is thought to contribute strongly to this effect. The latitude, mean temperature, daily maximum temperature and vapor pressure deficit also show significant associations, although these factors are correlated with each other in this dataset. Interestingly, stearic acid shows a very different correlation pattern than the other three fatty acids. The lack of significant correlations with water-related climate variables, such as the monthly climate moisture index, is also noteworthy. Further, the relationship between FAs was investigated. At first, the results of the GGE plot (Figure 2) may seem counter-intuitive. With the FA synthesis pathway in mind (Attia et al., 2021), an antagonistic relationship between oleic and linoleic acid is expected. However, one would expect the same kind of relationship between stearic and oleic acid, as they compete for the same precursor molecule. Instead, palmitic acid is more closely correlated with linoleic and oleic acid. However, once HO varieties are omitted, the resulting GGE biplot is in line with the expected result (Figure 2). In terms of a physiological trade-off, this suggests that the only way to increase oleic acid composition in HO genotypes once the stearic acid pool is depleted is to pull from palmitic acid further down the synthesis chain.

### 4.2 GWAS

The loci associated with mean FA content were consistent between environments when *FAD2-1* mutants were retained. As this consistency widely disappeared after omission of high oleic varieties, it might be valid to conclude that breeding for high oleic compositions resulted in trait stability and/or stability was actively selected for during breeding. It is also valid to conclude that the high oleic acid phenotype involves selection of loci beyond *FAD2-1*, a result that has been previously reported (Fernandez-Martinez et al., 1989; Lacombe et al., 2001). This hypothesis is in line with the results depicted in Figure 3, showing significant differences in both oleic and linoleic acid composition β stability between varieties with and without the *FAD2-1* mutation, as well as GWAS results. Unfortunately, not all 287 varieties of the SAM population were successfully grown under each condition. The largest number of varieties was 276 grown in Iowa 2014, the smallest in Iowa 2010 with 135. Only 45 were successfully phenotyped in all trials. This, in turn, means that the populations investigated vary, possibly influencing the differences between results from different trials beyond differences in climate, and negatively affecting power.

### 4.3 CANDIDATE GENE ANALYSIS

While a list of 429 genes involved in the FA metabolism is available (Badouin et al., 2017), this list was not exclusively used to subset the candidate genes in the hopes of capturing regulatory genes as well. This approach was successful, as in addition to promising FA metabolism genes such as oleosin and omega-3/omega-6 fatty acid desaturase, other classes such as heat shock proteins and transcription factors were found. Even an siRNA biogenesis regulator complex gene was associated with mean FA and β stability, which may be important for accumulation and stability of oleic acid accumulation given the nature of the *FAD2-1* mutation (Schuppert et al., 2006; Tschopp et al., 2017). The number of candidate genes not overlapping between α and β stability GWAS is noteworthy. It suggests that trait stability within and between environments might have quite different genetic bases, which may make sense as β stability is due to genotype-by-environment interaction and α stability is mostly driven by reversion of some high oleic lines to wild type fatty acid composition. Notably, *FAD2-1*, which is thought to play a major role in the high oleic genotypes is absent from the candidate list and more than 1 Mb away from the nearest significant SNP. The reason for this might lie in the presence of regions of low diversity and extended linkage disequilibrium in the genomes of *FAD2-1* mutant varieties, the result of known rapid truncation selection due to preferential breeding for high oleic phenotypes in recent decades. It is also conceivable that additional variation in this region of the genome is needed for the phenotype to emerge, making only SNPs associated with the *FAD2-1* duplication, as well as other variants, significant. This, in addition to the thresholds used for the identification of candidates, would favor an approach incorporating LD, haploblocks and introgressions into the pipeline to identify multigenic regions associated with FA phenotypes. In this vein, chromosome 5 in the Georgia 2010 trial in Figure 5 is noteworthy. Two of the significant peaks at 145 and 185 Mbp, one of which is also significant in the Minnesota early 2016 trial, lie just outside a known inversion/introgression spanning 148-177 Mb (Todesco et al., 2020), suggesting that the entire introgression is associated with individual seed homogeneity in relation to FA composition, in addition to the known effect on ligule width in *H. annuus* and seed size in *H. petiolaris* (Todesco et al., 2020), and pericarp strength in domesticated sunflower lines (White et al., 2025). Comparisons with candidate genes and regions from other papers proved difficult because of outdated technology. In the QTL analysis investigating fatty acid composition stability under water stress by Ebrahimi et al. (2008), associated marker positions on chromosomes were denominated in cM (Ebrahimi et al., 2008). Pérez-Vich et al. (2016) investigated high palmitic varieties using QTL mapping, identifying 3-ketoacyl-CoA synthase II (KCSII) as the causal gene (Pérez-Vich et al., 2016). In this work, we also identified a KCS gene as a candidate on chromosome 9, but as it is only supported by one SNP in one environment, its role in the present panel is uncertain. A prior GWAS on fatty acid composition by Chernova et al. (2021) identified several large LD blocks associated with oleic and linoleic acid, among other FAs not analyzed in the present study; however, none of these blocks appear to overlap with loci identified in the multivariate analyses of our study.

Fatty acid composition and stability are governed by the *FAD2-1* locus with significant effects at multiple other loci across the genome. Plasticity across environments (β stability) had fairly simple inheritance, which overlapped with loci involved in mean fatty acid composition. On the other hand, α stability, or the stability of the phenotype among plants within a genotype, varied among environments, but a few loci were important, including a region on chromosome 5 with a known structural variant that also influences other, unrelated traits. Overall, this work provides a basis to target loci in breeding to maintain stability of the fatty acid profile in sunflower, a key trait for this oilseed crop of worldwide importance.

## Acknowledgements

The authors would like to thank many volunteer and paid interns for assistance with the field studies, and especially Brady Koehler and Michael Grove of USDA-ARS for their invaluable coordination of the field work. Mention of trade names or commercial products in this article is solely for the purpose of providing specific information and does not imply recommendation or endorsement by the U.S. Department of Agriculture. USDA is an equal opportunity provider and employer.

## Funding

The authors wish to acknowledge funding from the U.S. Department of Agriculture—Agricultural Research Service CRIS projects 3060-21000-043-00D and 3060-21000-047-00D, a grant from the National Institute of Food and Agriculture 2008-35300-19263, and Genome Canada and Genome BC’s Applied Genomics Research in Bioproducts or Crops (ABC) Competition. McNellie is supported by a U.S. Department of Agriculture – Agricultural Research Service Postdoctoral Fellowship.

## Contributions

BSH, JMB, and LHR conceived of the research and sourced funding. QG, JRM, JGB, and BSH collected field samples. QG collected laboratory data. MI, QG, JPM, and KGK performed statistical modelling of genotype data and genotype-phenotype associations, MI and BSH wrote the manuscript with input from other authors. All authors have read and approved the final manuscript.

## Conflict of Interest

BSH serves on the editorial board for Theoretical and Applied Genetics.

## Notes

### Competing Interest Statement

The authors have declared no competing interest.

