## Supplemental Figure for "The stability of fatty acid composition in sunflower oil is dependent on environment and affected by structural variation"

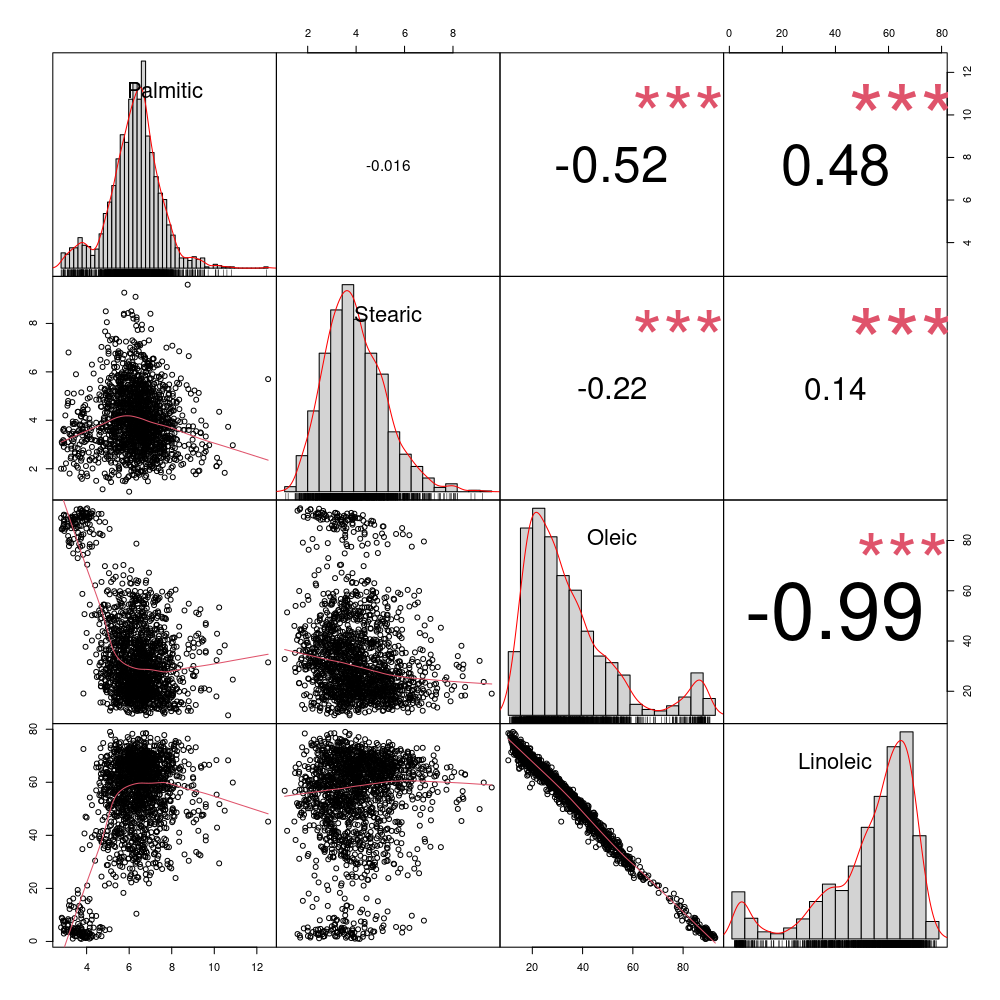

**Supplemental Figure 1: Fatty acid distributions across all samples in all environments.** Histograms of fatty acid percentages (diagonal), bivariate scatter plots with a fitted line (below diagonal) and the value of the correlation with significance level (***: p-value < 0.001), above diagonal.

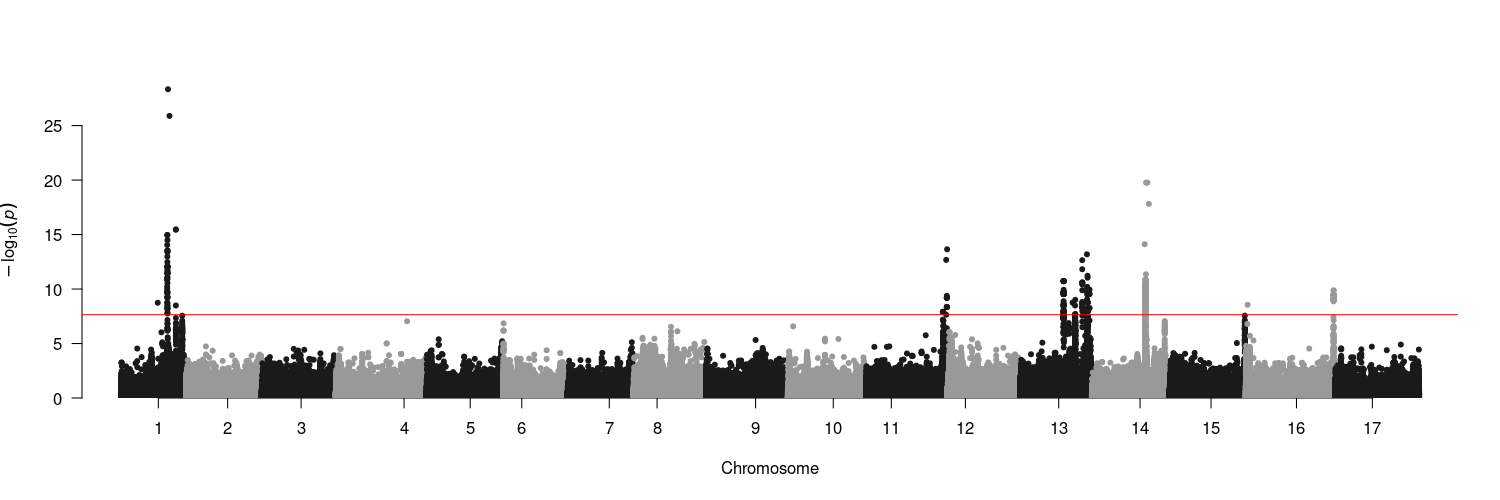

**Supplemental Figure 2:** **Manhattan plot of fatty acid stability across environments (Eberhart & Russell’s** β**).** SNP positions are depicted on the x-axis, -log10 of the p-value on the y-axis. The Bonferroni threshold based on the number of SNPs is depicted as a red line.

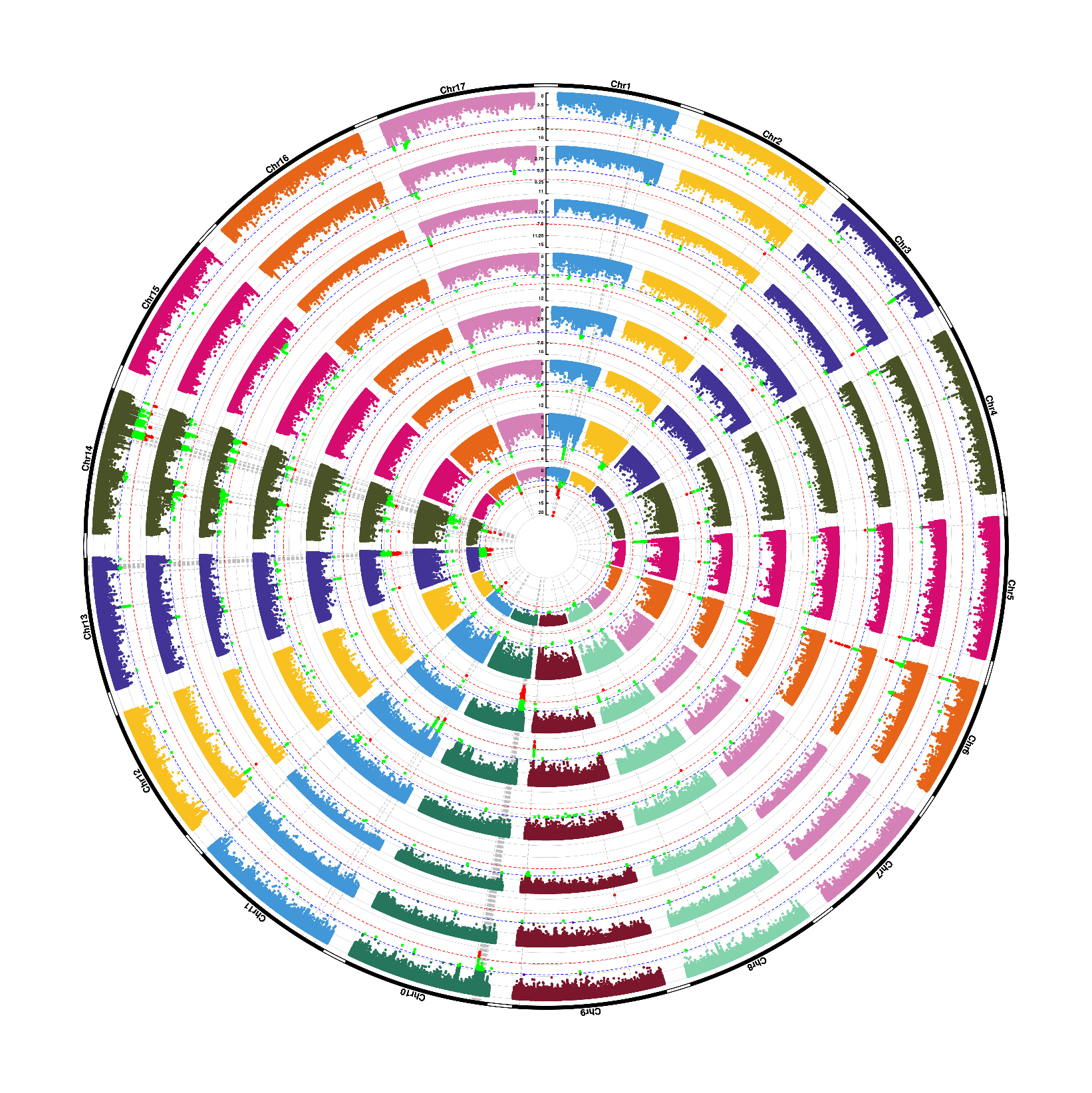

**Supplemental Figure 3: Manhattan plots of mean fatty acid compositions in all trials after omitting HO varieties.** From inside to outside, the Manhattan plots represent the results of the trials in British Columbia 2010, Minnesota 2015 and early and late 2016, Iowa 2010, 2013 and 2014 and Georgia 2010. SNP positions are depicted on the x-axis, -log10 of the p-value on the y-axis. The Bonferroni threshold based on the number of SNPs is depicted as a red line, the threshold based on the number of LD blocks as a blue line. SNPs passing the lower threshold are marked green, SNPs passing the more stringent threshold are marked in red.

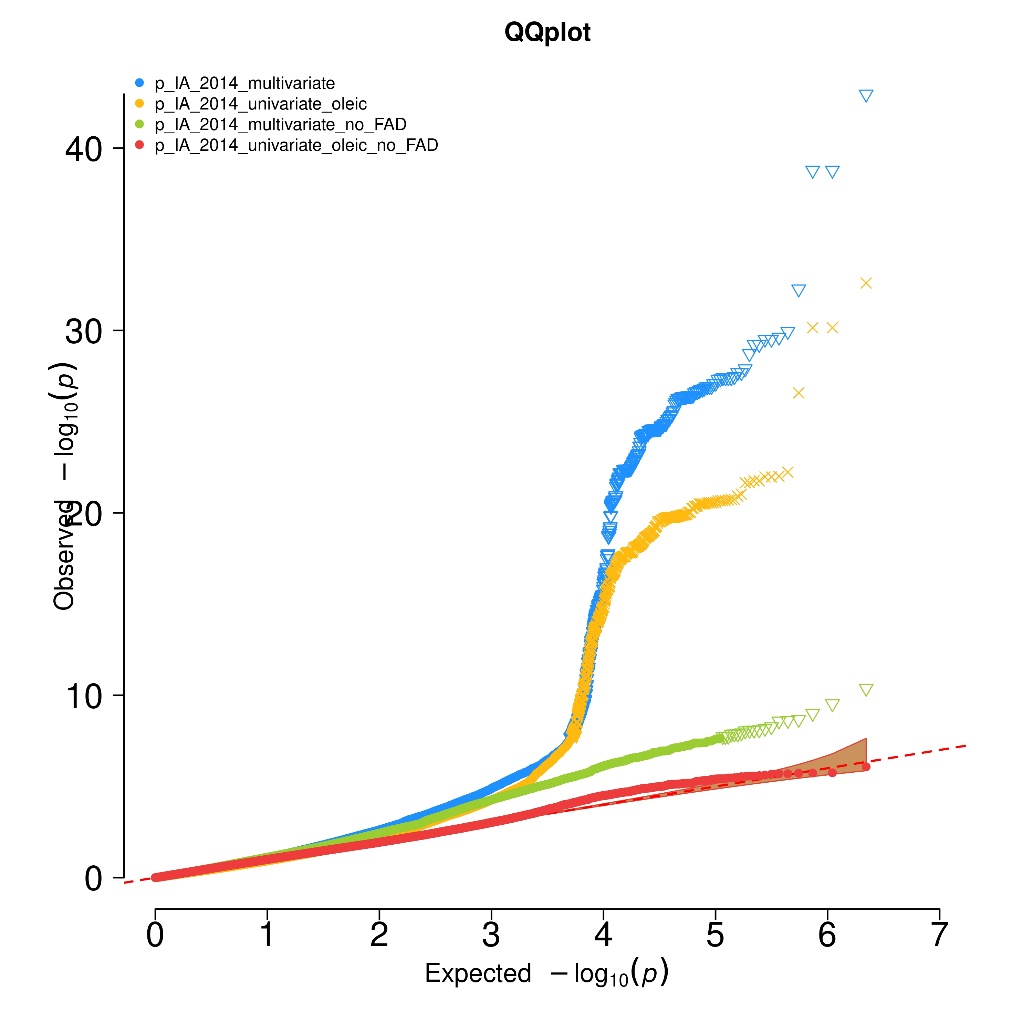

**Supplemental Figure 4:** **QQ plot comparing multivariate GWAS using all varieties vs. omission of high-oleic varieties.** Expected (x-axis) vs. observed (y-axis) p-values of multivariate GWAS on mean FA composition, with (blue) and without (green) HO varieties in the Iowa 2014 trial. Results for univariate GWAS on oleic acid with HO varieties (yellow) and with FAD2-1 mutation as a covariate (red) are shown as well. Crosses and triangles signify p-values below the Bonferroni threshold.

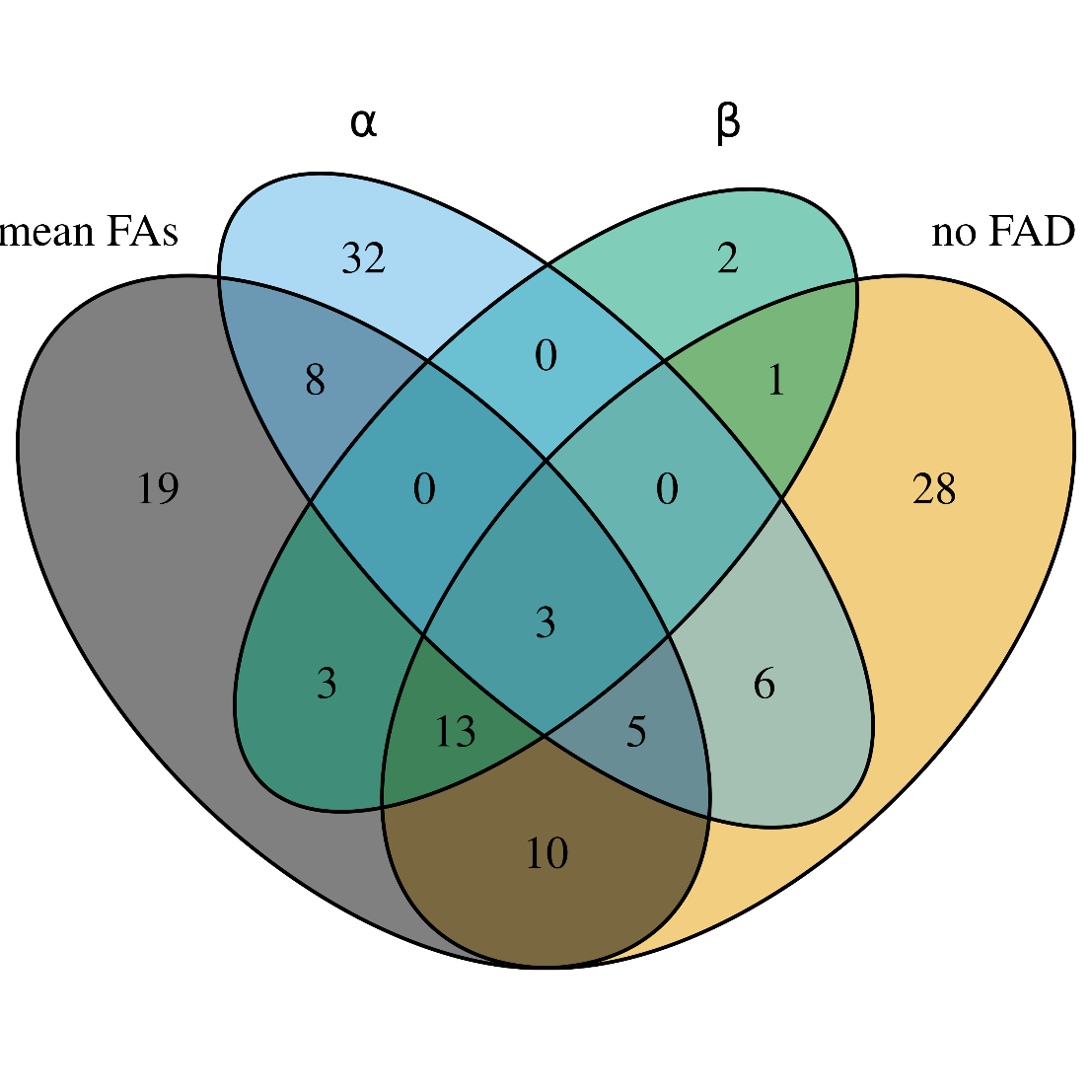

**Supplemental Figure 5: Venn Diagram of candidate genes identified in the four multivariate GWAS:** fatty acid composition without FAD mutant omission (mean FAs), with FAD mutant omission (no FAD), within-environment variation (α) and between-environment variation (β).

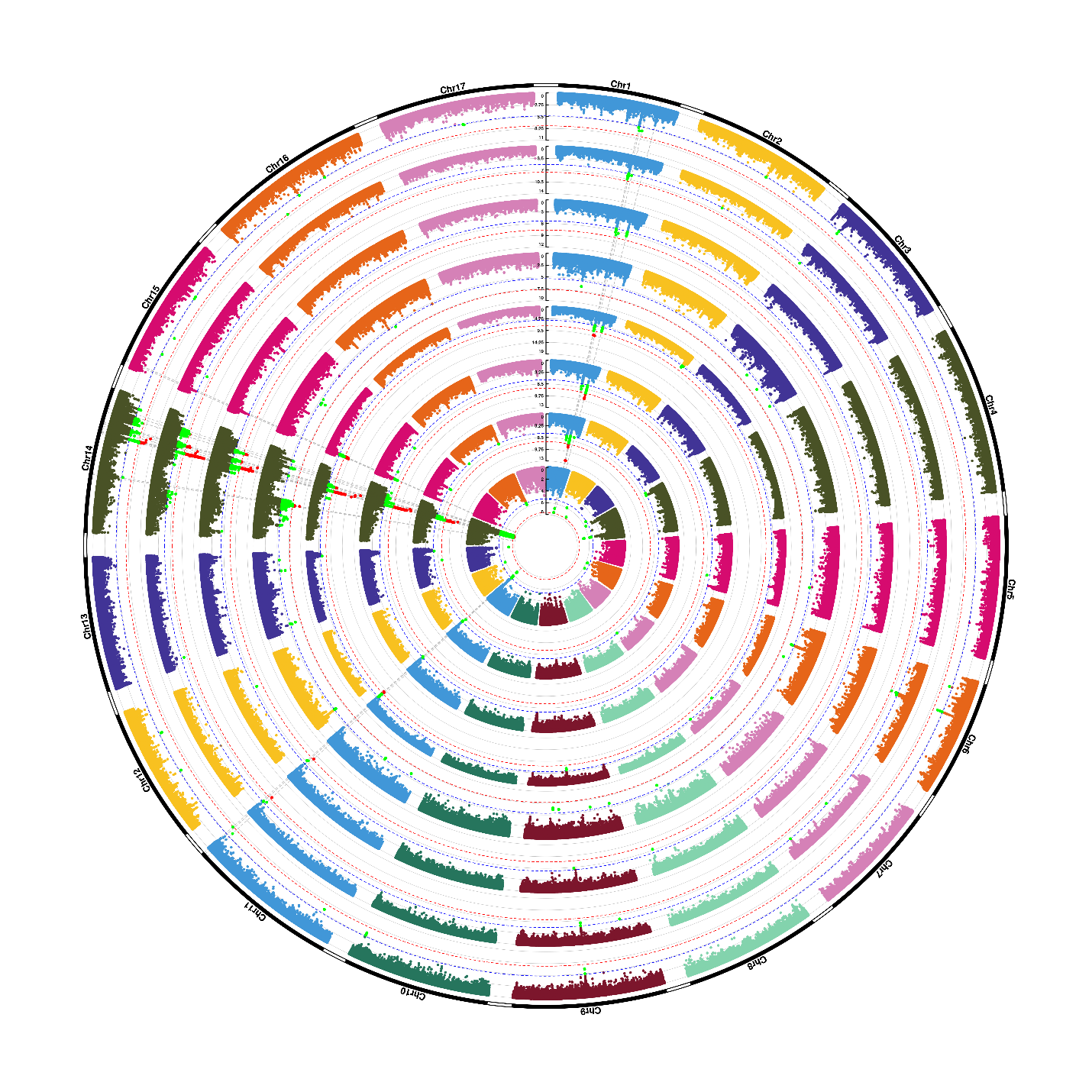

**Supplemental Figure 6:** **Manhattan plots of univariate tests of mean palmitic acid compositions in all trials.** From inside to outside, the Manhattan plots represent the results of the trials in British Columbia 2010, Minnesota 2015 and early and late 2016, Iowa 2010, 2013 and 2014 and Georgia 2010. SNP positions are depicted on the x-axis, -log10 of the p-value on the y-axis. The Bonferroni threshold based on the number of SNPs is depicted as a red line, the threshold based on the number of LD blocks as a blue line. SNPs passing the lower threshold are marked green, SNPs passing the more stringent threshold are marked in red.

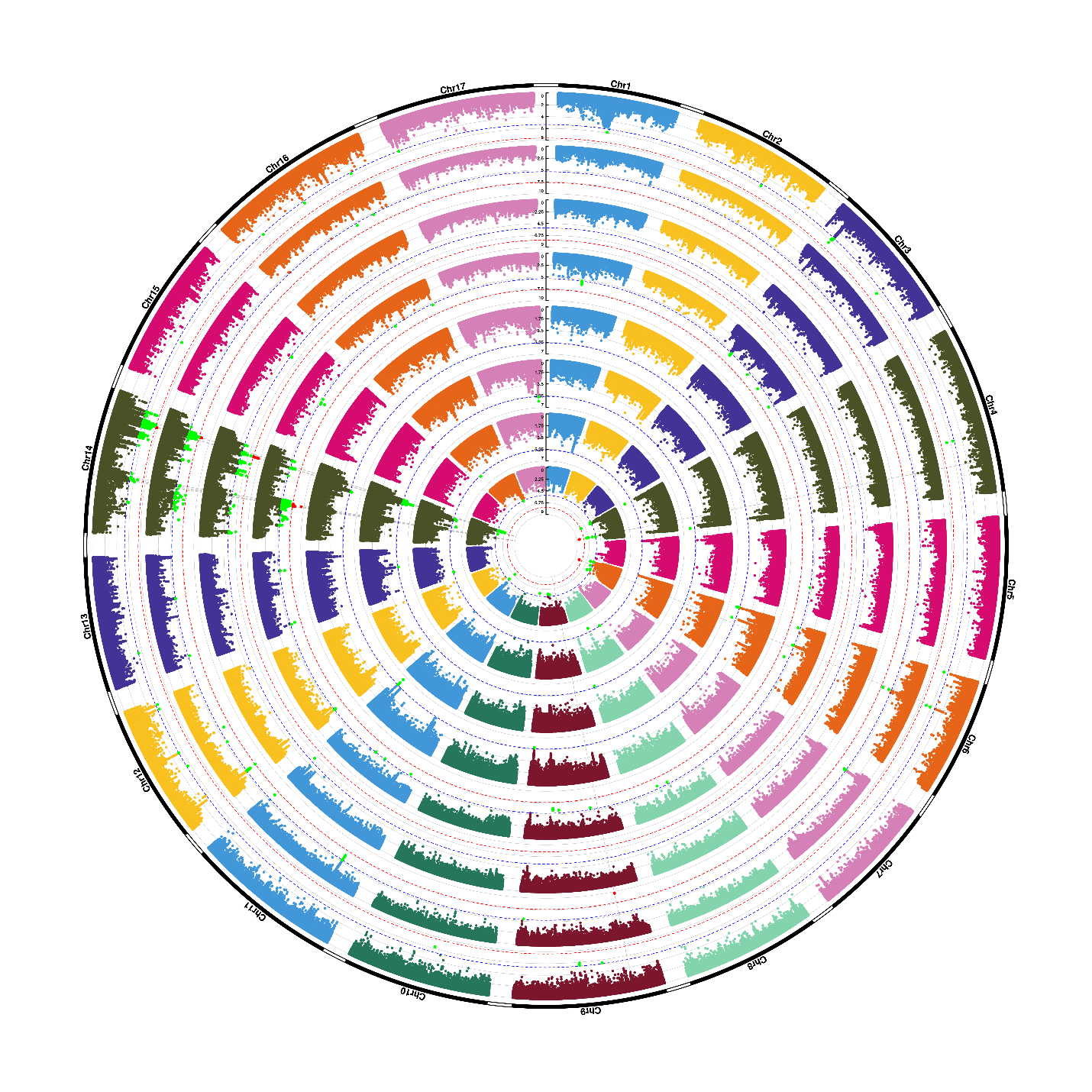
 **Supplemental Figure 7:** **Manhattan plots of univariate tests of mean palmitic acid compositions in all trials after adding the *FAD2-1* mutation as a covariate.** From inside to outside, the Manhattan plots represent the results of the trials in British Columbia 2010, Minnesota 2015 and early and late 2016, Iowa 2010, 2013 and 2014 and Georgia 2010. SNP positions are depicted on the x-axis, -log10 of the p-value on the y-axis. The Bonferroni threshold based on the number of SNPs is depicted as a red line, the threshold based on the number of LD blocks as a blue line. SNPs passing the lower threshold are marked green, SNPs passing the more stringent threshold are marked in red.

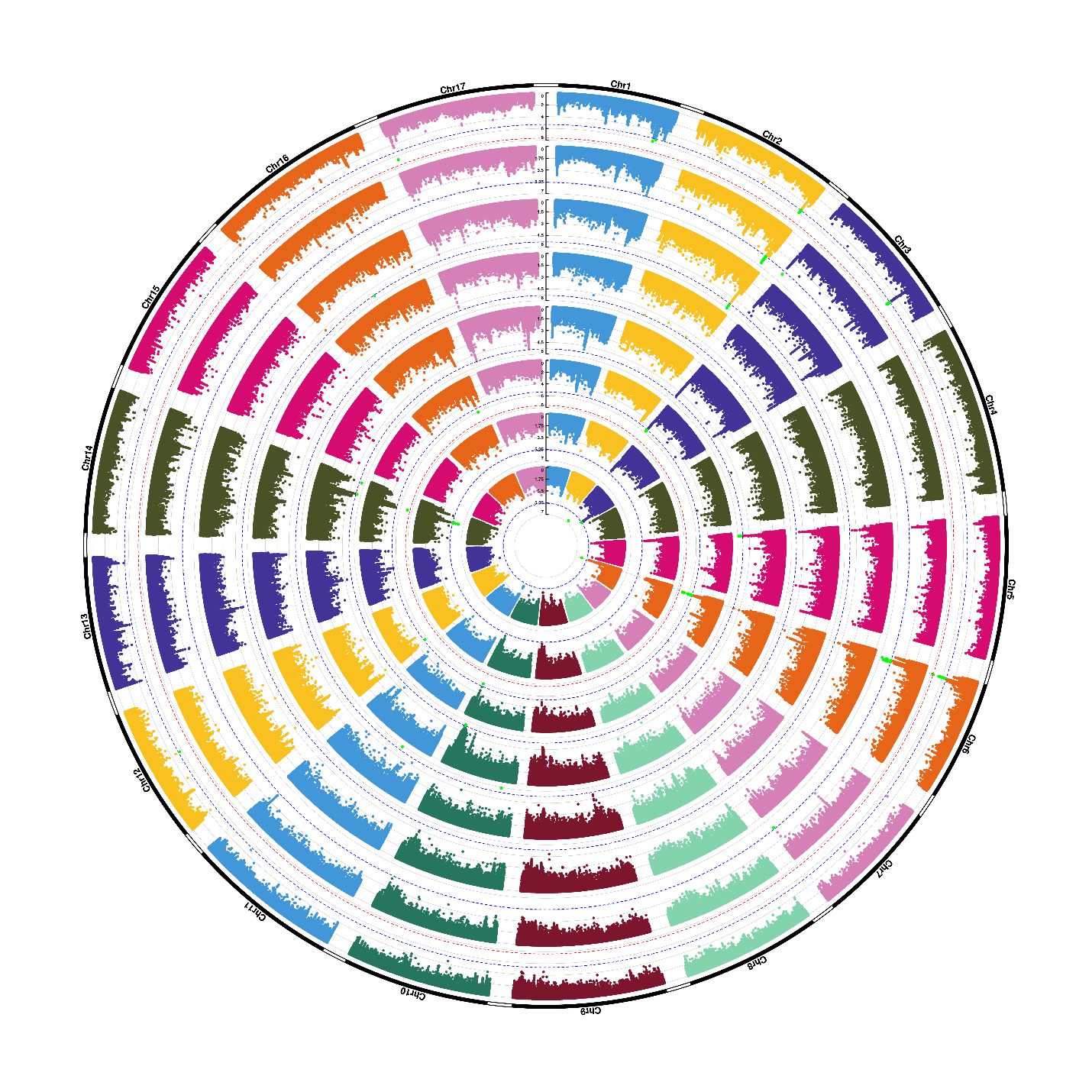
 **Supplemental Figure 8:** **Manhattan plots of univariate tests of mean stearic acid compositions in all trials.** From inside to outside, the Manhattan plots represent the results of the trials in British Columbia 2010, Minnesota 2015 and early and late 2016, Iowa 2010, 2013 and 2014 and Georgia 2010. SNP positions are depicted on the x-axis, -log10 of the p-value on the y-axis. The Bonferroni threshold based on the number of SNPs is depicted as a red line, the threshold based on the number of LD blocks as a blue line. SNPs passing the lower threshold are marked green, SNPs passing the more stringent threshold are marked in red.

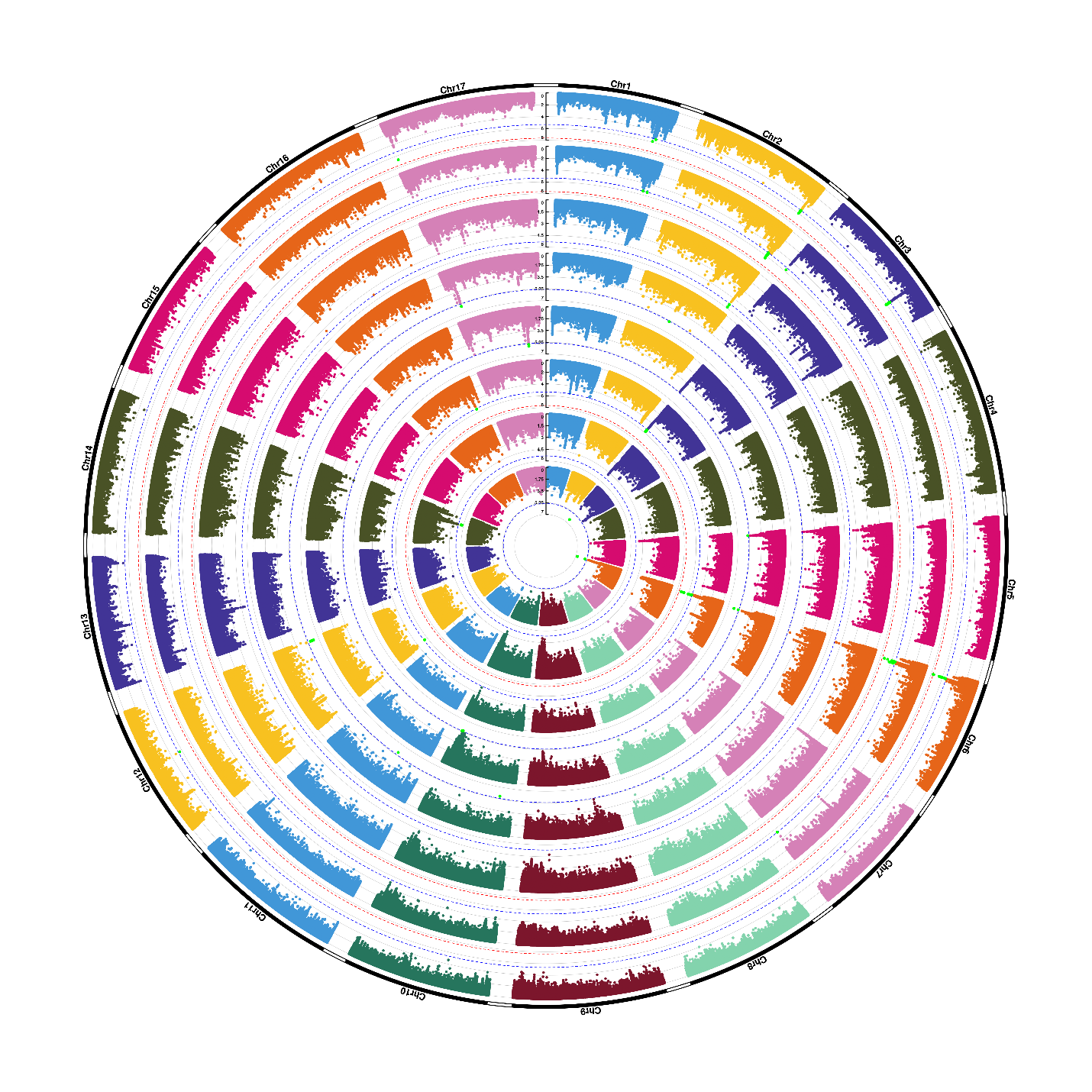
 **Supplemental Figure 9:** **Manhattan plots of univariate tests of mean stearic acid compositions in all trials after adding the *FAD2-1* mutation as a covariate.** From inside to outside, the Manhattan plots represent the results of the trials in British Columbia 2010, Minnesota 2015 and early and late 2016, Iowa 2010, 2013 and 2014 and Georgia 2010. SNP positions are depicted on the x-axis, -log10 of the p-value on the y-axis. The Bonferroni threshold based on the number of SNPs is depicted as a red line, the threshold based on the number of LD blocks as a blue line. SNPs passing the lower threshold are marked green, SNPs passing the more stringent threshold are marked in red.

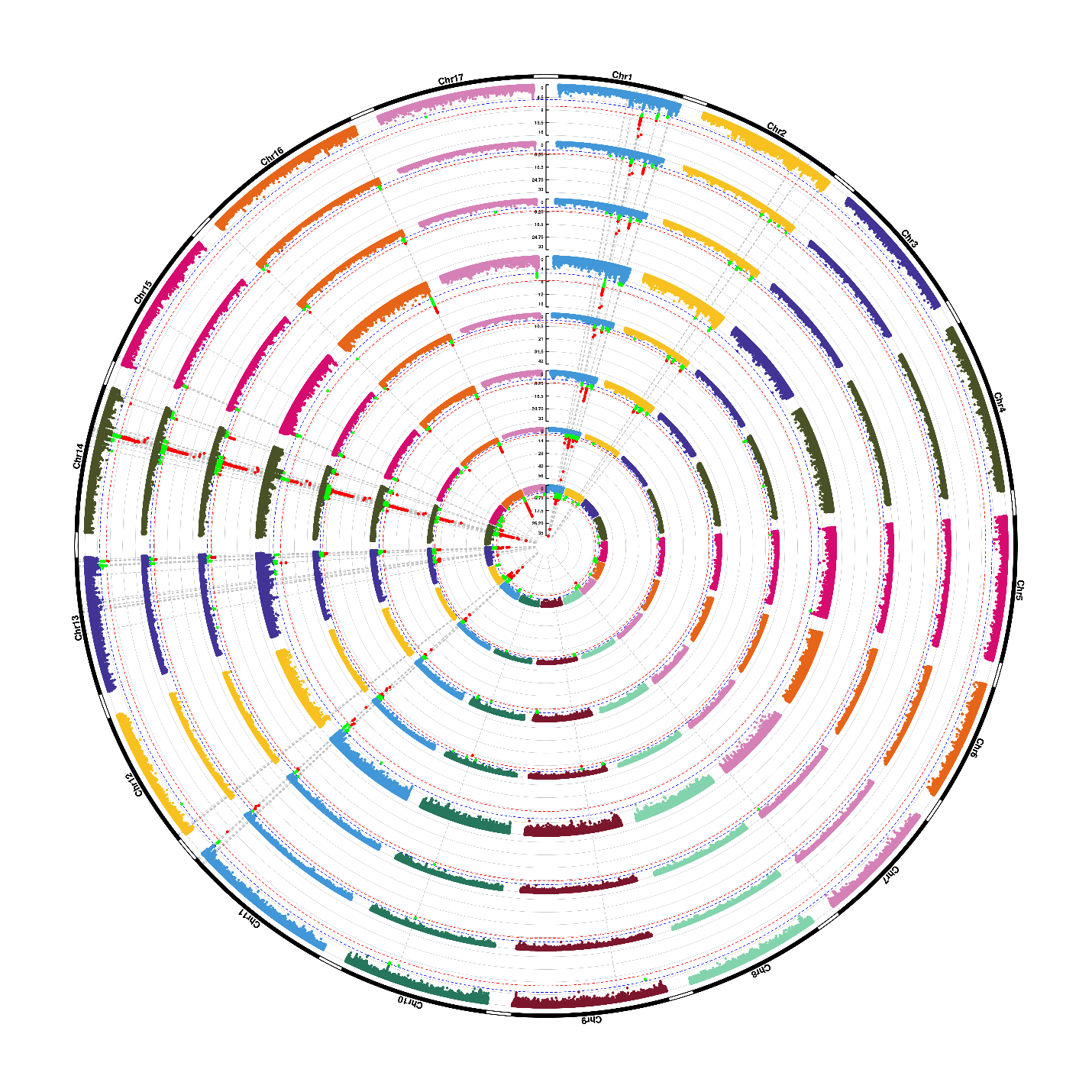

**Supplemental Figure 10:** **Manhattan plots of univariate tests of mean oleic acid compositions in all trials.** From inside to outside, the Manhattan plots represent the results of the trials in British Columbia 2010, Minnesota 2015 and early and late 2016, Iowa 2010, 2013 and 2014 and Georgia 2010. SNP positions are depicted on the x-axis, -log10 of the p-value on the y-axis. The Bonferroni threshold based on the number of SNPs is depicted as a red line, the threshold based on the number of LD blocks as a blue line. SNPs passing the lower threshold are marked green, SNPs passing the more stringent threshold are marked in red.

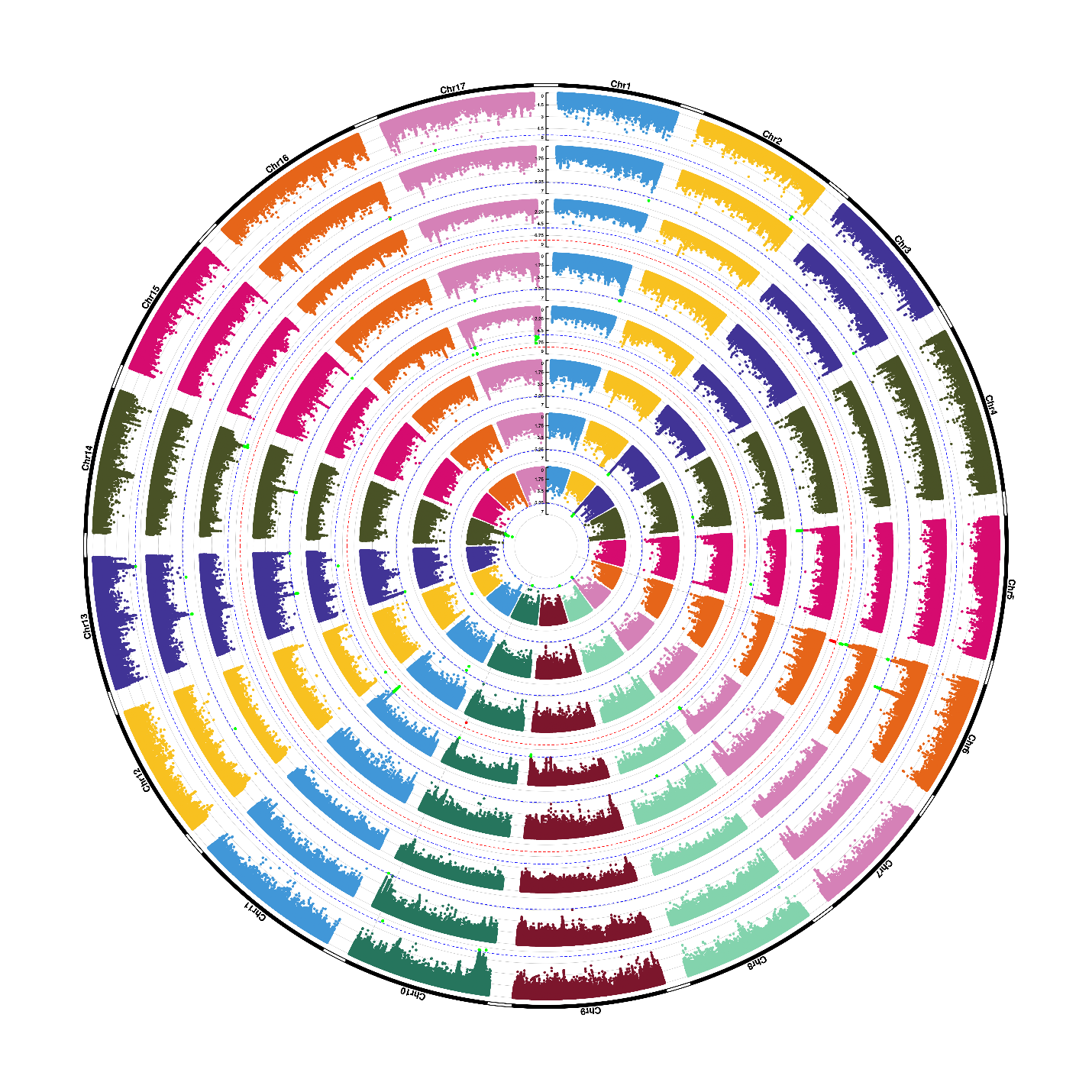
 **Supplemental Figure 11:** **Manhattan plots of univariate tests of mean oleic acid compositions in all trials after adding the *FAD2-1* mutation as a covariate.** From inside to outside, the Manhattan plots represent the results of the trials in British Columbia 2010, Minnesota 2015 and early and late 2016, Iowa 2010, 2013 and 2014 and Georgia 2010. SNP positions are depicted on the x-axis, -log10 of the p-value on the y-axis. The Bonferroni threshold based on the number of SNPs is depicted as a red line, the threshold based on the number of LD blocks as a blue line. SNPs passing the lower threshold are marked green, SNPs passing the more stringent threshold are marked in red.

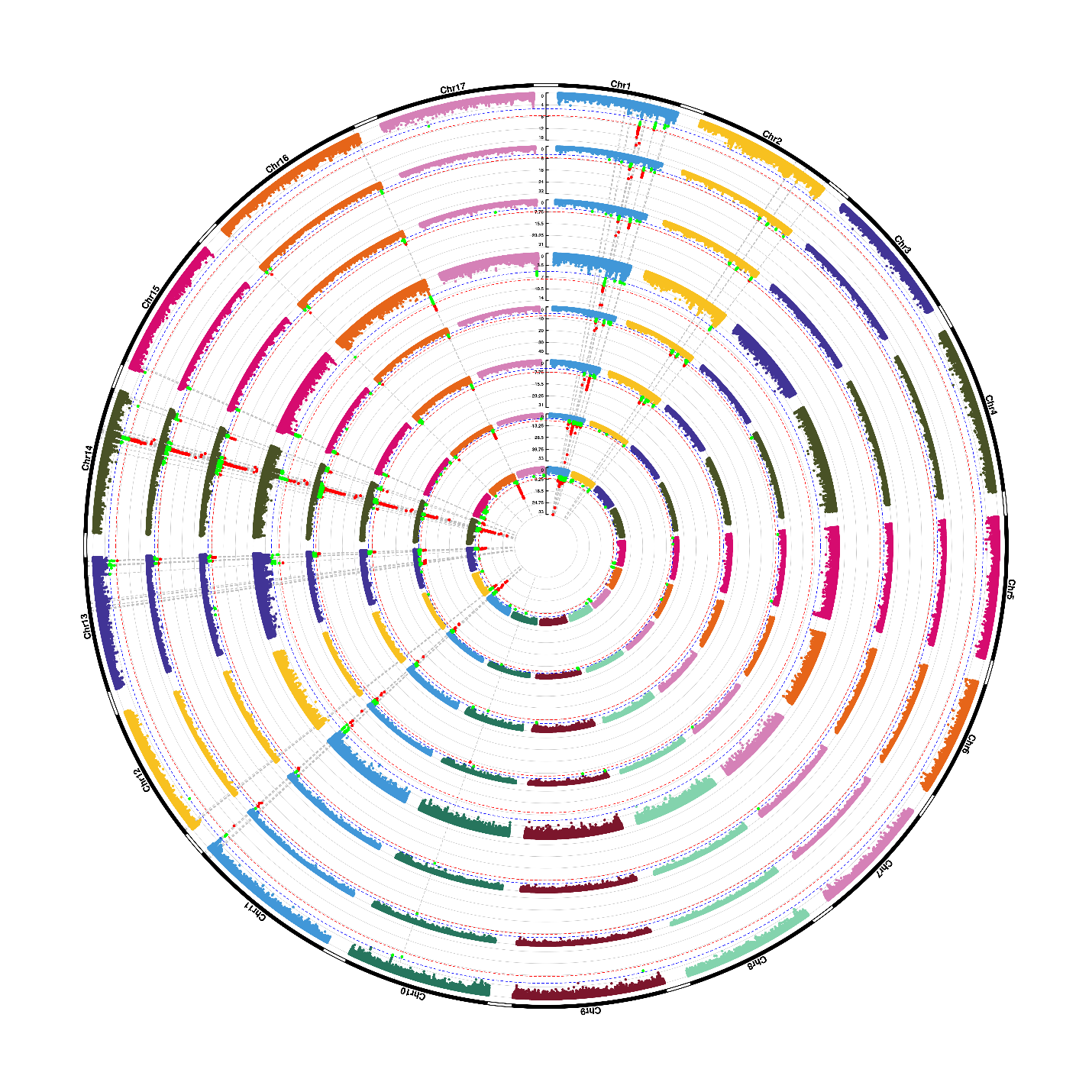
 **Supplemental Figure 12: Manhattan plots of univariate tests of mean linoleic acid compositions in all trials.** From inside to outside, the Manhattan plots represent the results of the trials in British Columbia 2010, Minnesota 2015 and early and late 2016, Iowa 2010, 2013 and 2014 and Georgia 2010. SNP positions are depicted on the x-axis, -log10 of the p-value on the y-axis. The Bonferroni threshold based on the number of SNPs is depicted as a red line, the threshold based on the number of LD blocks as a blue line. SNPs passing the lower threshold are marked green, SNPs passing the more stringent threshold are marked in red.

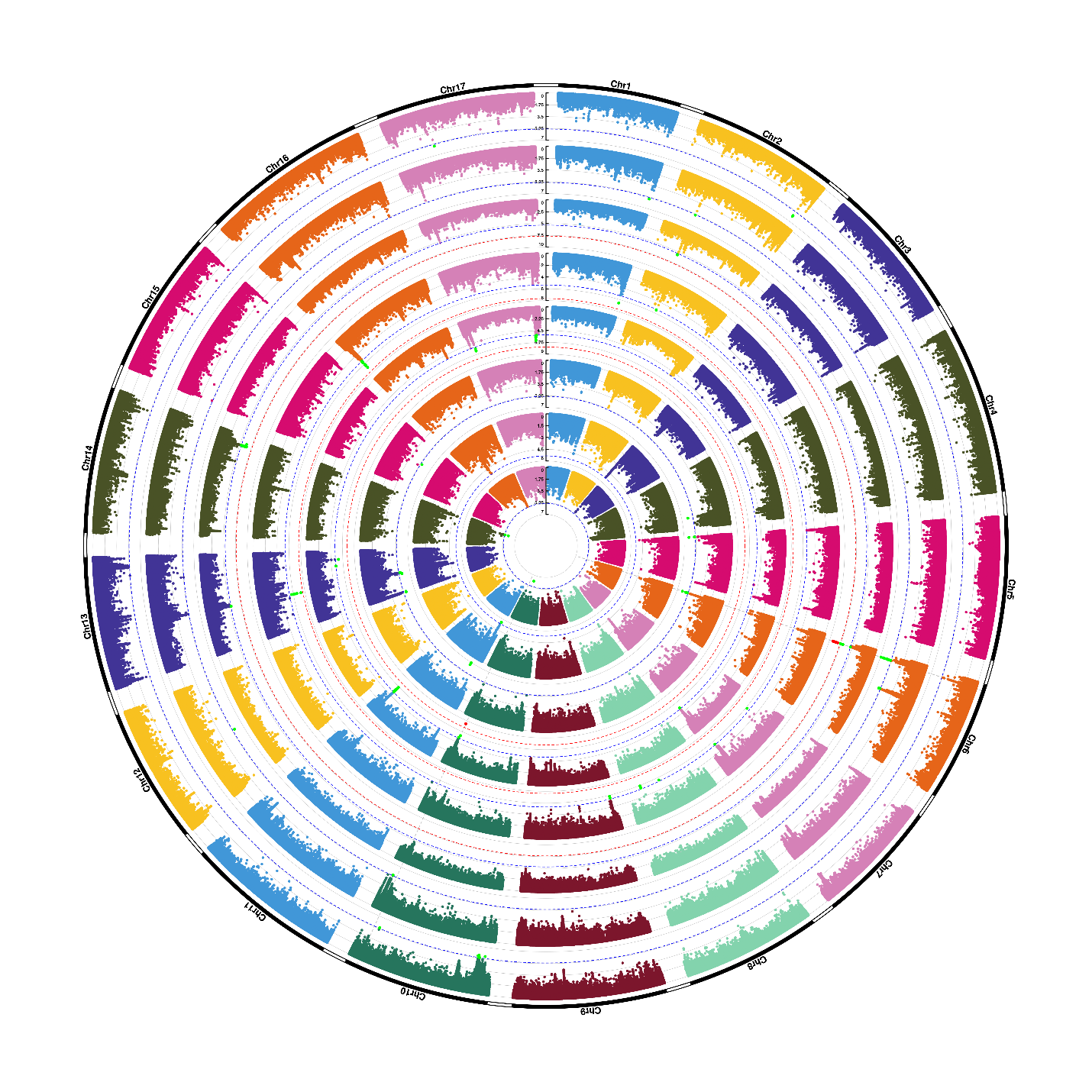

**Supplemental Figure 13:** **Manhattan plots of univariate tests of mean linoleic acid compositions in all trials after adding the *FAD2-1* mutation as a covariate.** From inside to outside, the Manhattan plots represent the results of the trials in British Columbia 2010, Minnesota 2015 and early and late 2016, Iowa 2010, 2013 and 2014 and Georgia 2010. SNP positions are depicted on the x-axis, -log10 of the p-value on the y-axis. The Bonferroni threshold based on the number of SNPs is depicted as a red line, the threshold based on the number of LD blocks as a blue line. SNPs passing the lower threshold are marked green, SNPs passing the more stringent threshold are marked in red.

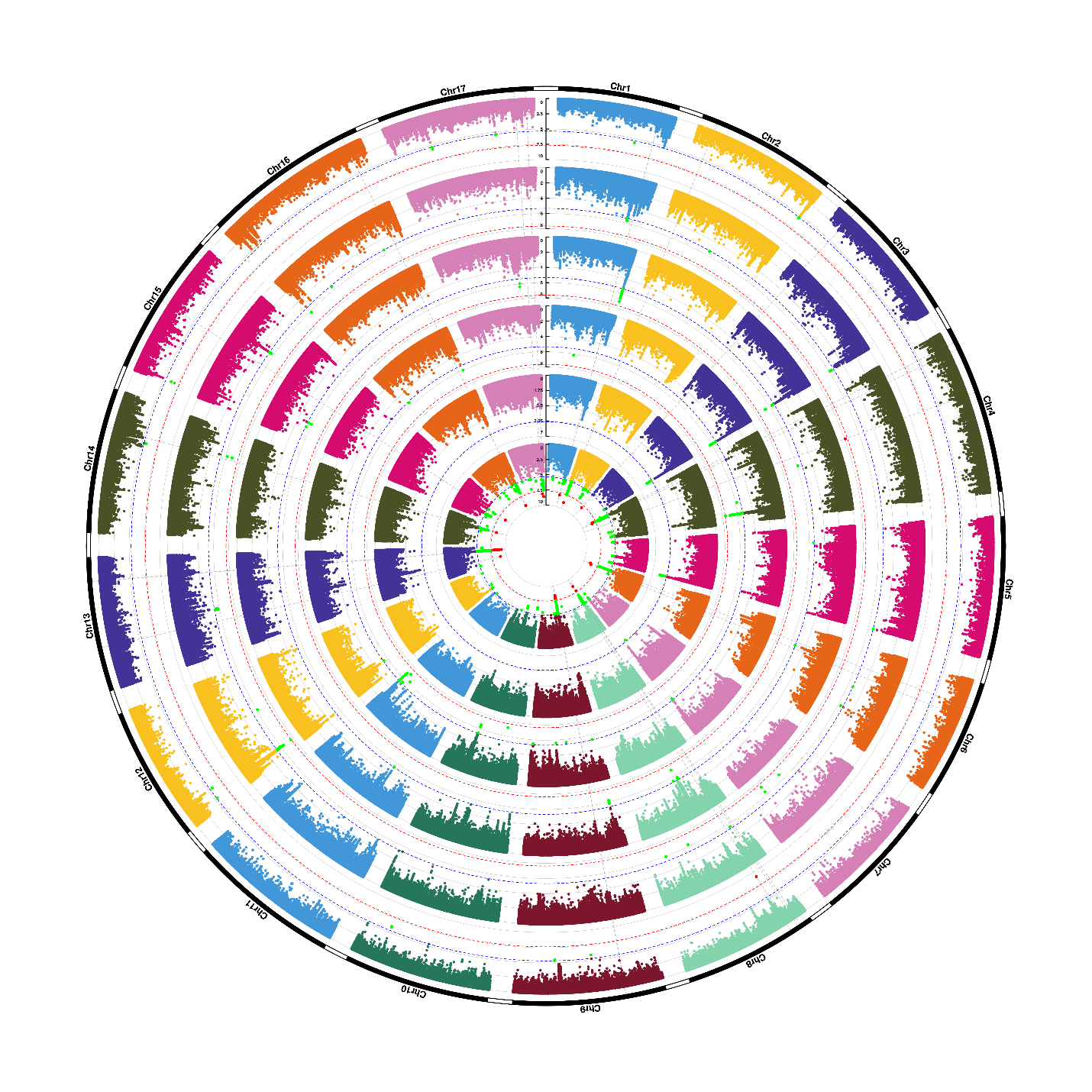
 **Supplemental Figure 14:** **Manhattan plots of univariate tests of mean palmitic acid α-stability.** From inside to outside, the Manhattan plots represent the results of the trials in British Columbia 2010, Minnesota early and late 2016, Iowa 2013 and 2014, and Georgia 2010. SNP positions are depicted on the x-axis, -log10 of the p-value on the y-axis. The Bonferroni threshold based on the number of SNPs is depicted as a red line, the threshold based on the number of LD blocks as a blue line. SNPs passing the lower threshold are marked green, SNPs passing the more stringent threshold are marked in red.

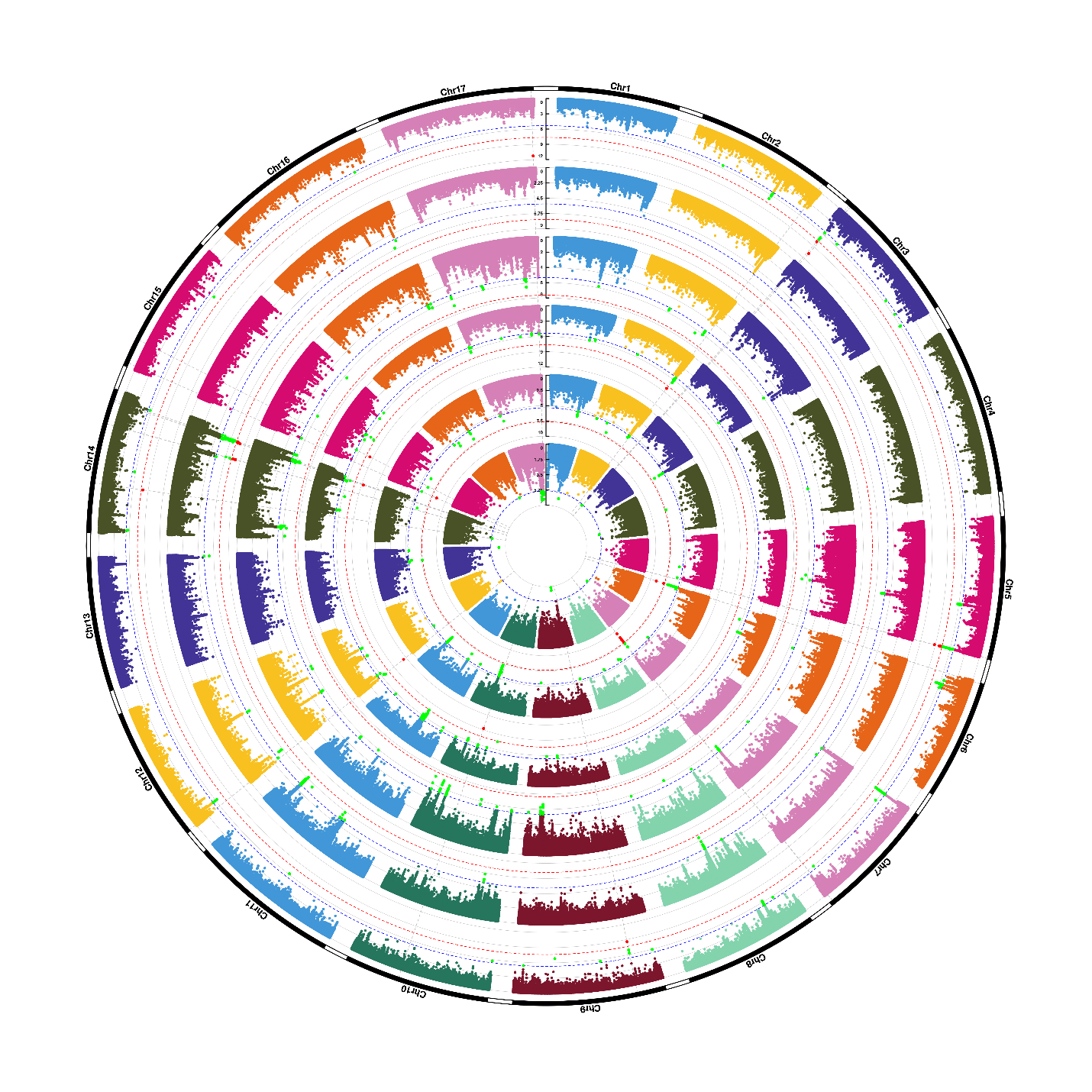
 **Supplemental Figure 15:** **Manhattan plots of univariate tests of mean stearic acid α-stability.** From inside to outside, the Manhattan plots represent the results of the trials in British Columbia 2010, Minnesota early and late 2016, Iowa 2013 and 2014, and Georgia 2010. SNP positions are depicted on the x-axis, -log10 of the p-value on the y-axis. The Bonferroni threshold based on the number of SNPs is depicted as a red line, the threshold based on the number of LD blocks as a blue line. SNPs passing the lower threshold are marked green, SNPs passing the more stringent threshold are marked in red.

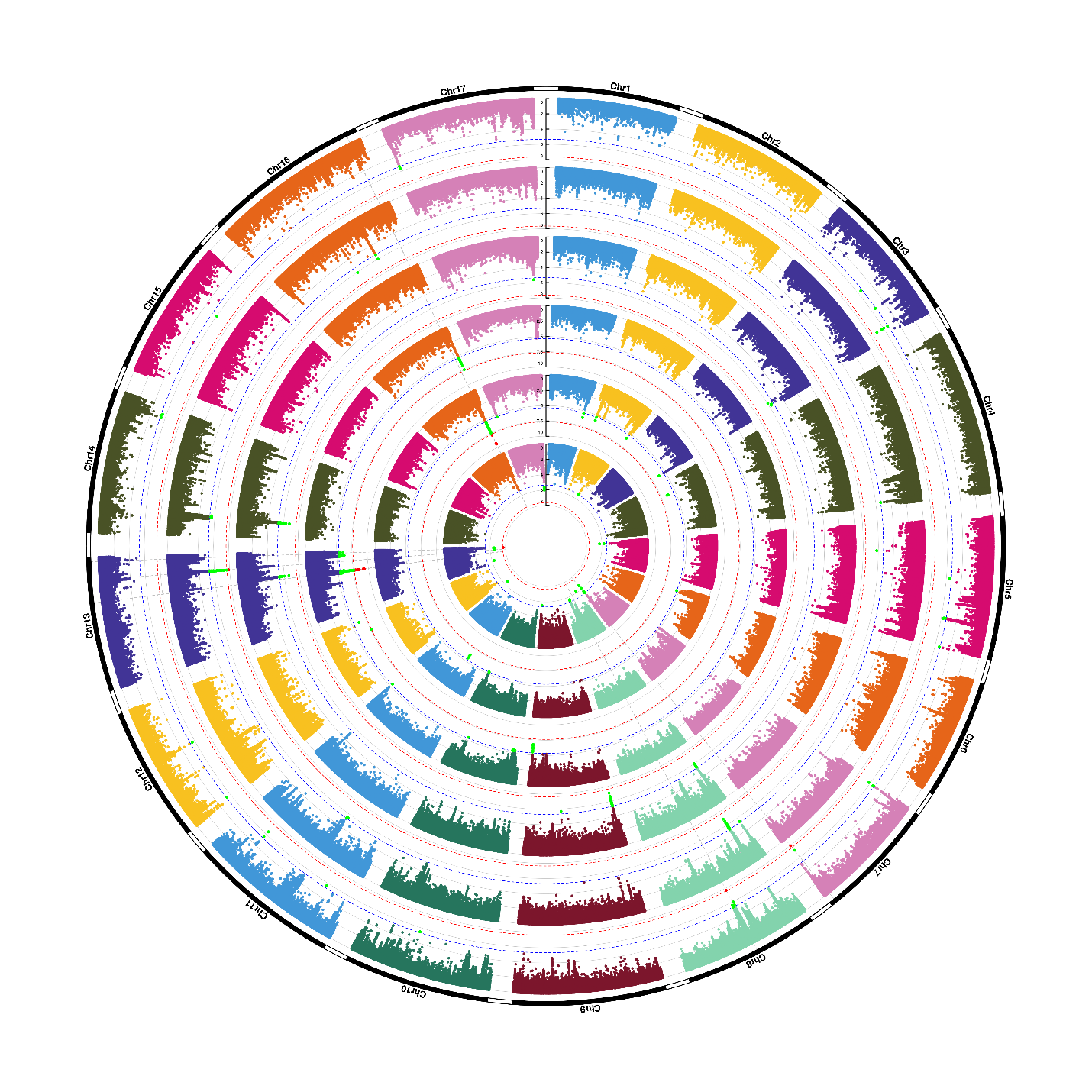
 **Supplemental Figure 16:** **Manhattan plots of univariate tests of mean oleic acid α-stability.** From inside to outside, the Manhattan plots represent the results of the trials in British Columbia 2010, Minnesota early and late 2016, Iowa 2013 and 2014, and Georgia 2010. SNP positions are depicted on the x-axis, -log10 of the p-value on the y-axis. The Bonferroni threshold based on the number of SNPs is depicted as a red line, the threshold based on the number of LD blocks as a blue line. SNPs passing the lower threshold are marked green, SNPs passing the more stringent threshold are marked in red.

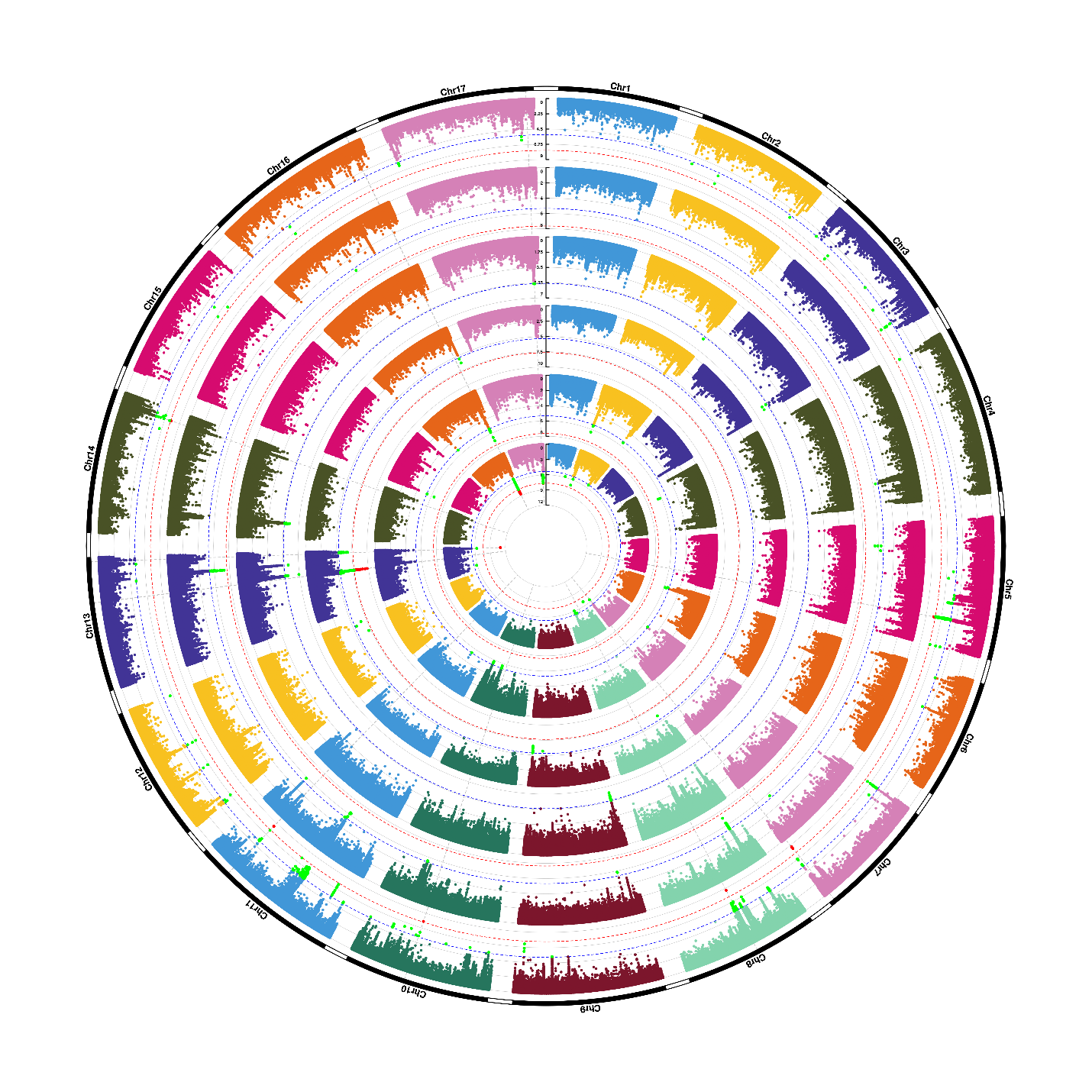
 **Supplemental Figure 17:** **Manhattan plots of univariate tests of mean linoleic acid α-stability.** From inside to outside, the Manhattan plots represent the results of the trials in British Columbia 2010, Minnesota early and late 2016, Iowa 2013 and 2014, and Georgia 2010. SNP positions are depicted on the x-axis, -log10 of the p-value on the y-axis. The Bonferroni threshold based on the number of SNPs is depicted as a red line, the threshold based on the number of LD blocks as a blue line. SNPs passing the lower threshold are marked green, SNPs passing the more stringent threshold are marked in red.
